# Parvalbumin interneuron ErbB4 controls ongoing network oscillations and olfactory behaviours in mice

**DOI:** 10.1101/2024.07.25.604407

**Authors:** Bin Hu, Chi Geng, Feng Guo, Ying Liu, Ran Wang, You-Ting Chen, Xiao-Yu Hou

**Author notes:** These authors contributed equally to this work (Bin Hu, Chi Geng). To whom correspondence may be addressed. (X.-Y. Hou).

## Abstract

Parvalbumin (PV)-positive interneurons modulate the processing of odour information. However, little is known about how PV interneurons dynamically remodel neural circuit responses in the olfactory bulb (OB) and their physiological significance. This study showed that a reinforced odour-discrimination task upregulated ErbB4 kinase activity in the mouse OB. ErbB4 knockout in the OB impaired the dishabituation of odour responses and discrimination of complex odours, whereas odour memory or adaptation was not altered in mice. ErbB4-positive neurones were localised throughout the OB, whereas ErbB4 was largely expressed in PV-positive interneurons within the internal and external plexiform layers. Similarly, ErbB4 ablation in PV interneurons disrupted olfactory discrimination and dishabituation in mice. ErbB4 knockout in PV interneurons disrupted the odour-evoked responses of mitral/tufted cells and increased the power of the ongoing local field potential in awake mice. We also observed a decrease in the frequency of miniature inhibitory postsynaptic currents and deficits in stimulus-evoked recurrent and lateral inhibition of mitral cells following PV-ErbB4 loss, suggesting a broad impairment in inhibitory microcircuits. These findings demonstrate that PV-ErbB4 signalling regulates inhibitory microcircuit activity, ongoing oscillations, and OB output, which underlie normal olfactory behaviour.

## Introduction

Discriminating chemical cues in the environment is not only necessary for evaluating potential threats, but also influences cognitive and emotional behaviours. The olfactory system dynamically adjusts its sensitivity to odours in response to changes in environmental, motivational, and cognitive states. For example, odour learning improves the performance of olfactory discrimination tests. Habituation and adaptation to successive exposure to an odour lead to decreased behavioural responses, whereas an unexpected odour enhances olfactory salience and stimulates behavioural dishabituation. Dysfunction in olfactory discrimination has been observed in the early stages of several neurodegenerative disorders and psychiatric diseases (Carnemolla et al., 2020; Walker et al., 2021). Similarly, the olfactory discrimination ability has a significant predictive value for future cognitive decline in healthy ageing individuals (Sohrabi et al., 2012; Uchida et al., 2020; Yoshitake et al., 2022). However, the neural mechanisms underlying the dynamic modulation of olfactory behaviour remain largely unknown.

The olfactory bulb (OB) is the first relay and processing station in the olfactory system. In the OB, odour signals are transformed into informational outputs that are sent to the associated olfactory cortex. As the principal output neurones of the OB, mitral/tufted cells (M/TCs) receive excitatory glutamatergic inputs from olfactory sensory neurones and inhibitory GABAergic inputs from heterogeneous subpopulations of interneurons. An imbalance in the excitation/inhibition onto mitral cells (MCs) impairs olfactory discrimination (Lepousez and Lledo, 2013). Furthermore, overlapping odour-evoked input patterns in the OB undergo reformatting to enhance the discrimination of similar odours, which depends on GABAergic interneurons (Gschwend et al., 2015; Li et al., 2018; Mohamed et al., 2019; Zavitz et al., 2020). GABAergic interneurons in the OB mediate the recurrent and lateral inhibition of MCs. Recurrent inhibition of MCs by granule cells (GCs) increases the frequency of odour-induced activity and facilitates odour discrimination (Aghvami et al., 2022; MacLeod and Laurent, 1996; Nunes and Kuner, 2015). Interglomerular lateral inhibition sharpens or filters the odour-induced output signals of MCs, enhancing the contrast of odour representations and, consequently, the ability to discriminate between similar odours (Egger and Kuner, 2021). Overall, the OB contains various interneuron types that control different olfactory behaviours (Lyons-Warren et al., 2023; Pardasani et al., 2023; Takahashi et al., 2016). Parvalbumin-positive (PV) interneurons form dendrodendritic contacts with neighbouring MCs ( Huang et al., 2013; Kato et al., 2013; Matsuno et al., 2017; Miyamichi et al., 2013). Unlike the odour-specific responses of GCs and MCs, PV interneurons exhibit strong responses to a broader range of odours (Kato et al., 2013). M/TCs receive significant inputs from PV interneurons from the external plexiform layer (EPL) (Miyamichi et al., 2013). However, the contribution of PV interneurons to olfactory behaviour and the associated molecular mechanisms are not understood.

GABAergic neurones preferentially express the receptor tyrosine kinase ErbB4 in cortical areas; however, in subcortical regions, ErbB4 is also expressed by non-GABAergic neurones (Bean et al., 2014; Deng et al., 2019; Dominguez et al., 2019; Fazzari et al., 2010; Lin et al., 2018; Mei and Nave, 2014; Shi and Bergson, 2020). ErbB4 kinases are expressed in neuroblasts in the subventricular zone and rostral migratory stream to the OB, implicating ErbB4 signalling in olfactory interneuronal precursor differentiation and olfaction (Anton et al., 2004). ErbB4 regulates the synaptic transmission and plasticity of PV interneurons in the neocortex and hippocampus (Chen et al., 2022; Dominguez et al., 2019; Grieco et al., 2020; Lin et al., 2018). Recently, we found a distinct ErbB4 signalling pathway in PV interneurons of the prefrontal cortex that hinders flexibility in odour-associated memory updating but has no influence on initial olfactory learning and memory (Cai and Shuman, 2022; Xu et al., 2022). However, whether PV neurones express ErbB4 in the OB and whether this affects olfactory behaviour remain unclear.

In this study, we showed that an odour-discrimination task promotes the activation of ErbB4 in the OB. ErbB4-positive neurones are localised throughout the OB, with most being PV-positive interneurons in the internal plexiform layer (IPL) and EPL. In this context, we evaluated the performance of PV-ErbB4 knockout mice in odour discrimination, habituation/dishabituation, and detection threshold tasks. We also explored the effects of ErbB4 in PV interneurons on the odour-evoked activity of MCs and ongoing network oscillations in awake mice, as well as on the inhibitory modulation of MC output. Our findings reveal the physiological significance of PV interneurons via ErbB4 signalling in the dynamic modulation of olfactory behaviour.

## Results

### Performing an odour discrimination task upregulates the activity of ErbB4 in the OB

First, we examined the levels of ErbB4 protein and its phosphorylated form (p-ErbB4) in mouse OB after a reinforced go/no-go odour discrimination task. A go/go task without odour discrimination was performed as a control (Figure 1A). Adult mice were first trained over 3 days of go/no-go tasks (200 trials per day) to discriminate between a pair of simple odours (water-rewarded isoamyl acetate and unrewarded heptanone) and then to discriminate complex chemical cues (60%/40% binary mixtures of these odours, “6/4” refers to 60% isoamyl acetate + 40% heptanone, and “4/6” refers to 40% isoamyl acetate + 60% heptanone, in volume) (Figure 1B). As shown in Figure 1C and 1D (see Figures 1 and 2 and Supplementary Figure 1A and 1 B for uncropped western blot images), the level of p-ErbB4 in the go/no-go group was higher than that in the go/go group, implying a link between increased ErbB4 activity in the OB and olfactory discrimination ability.

**Figure 1.**
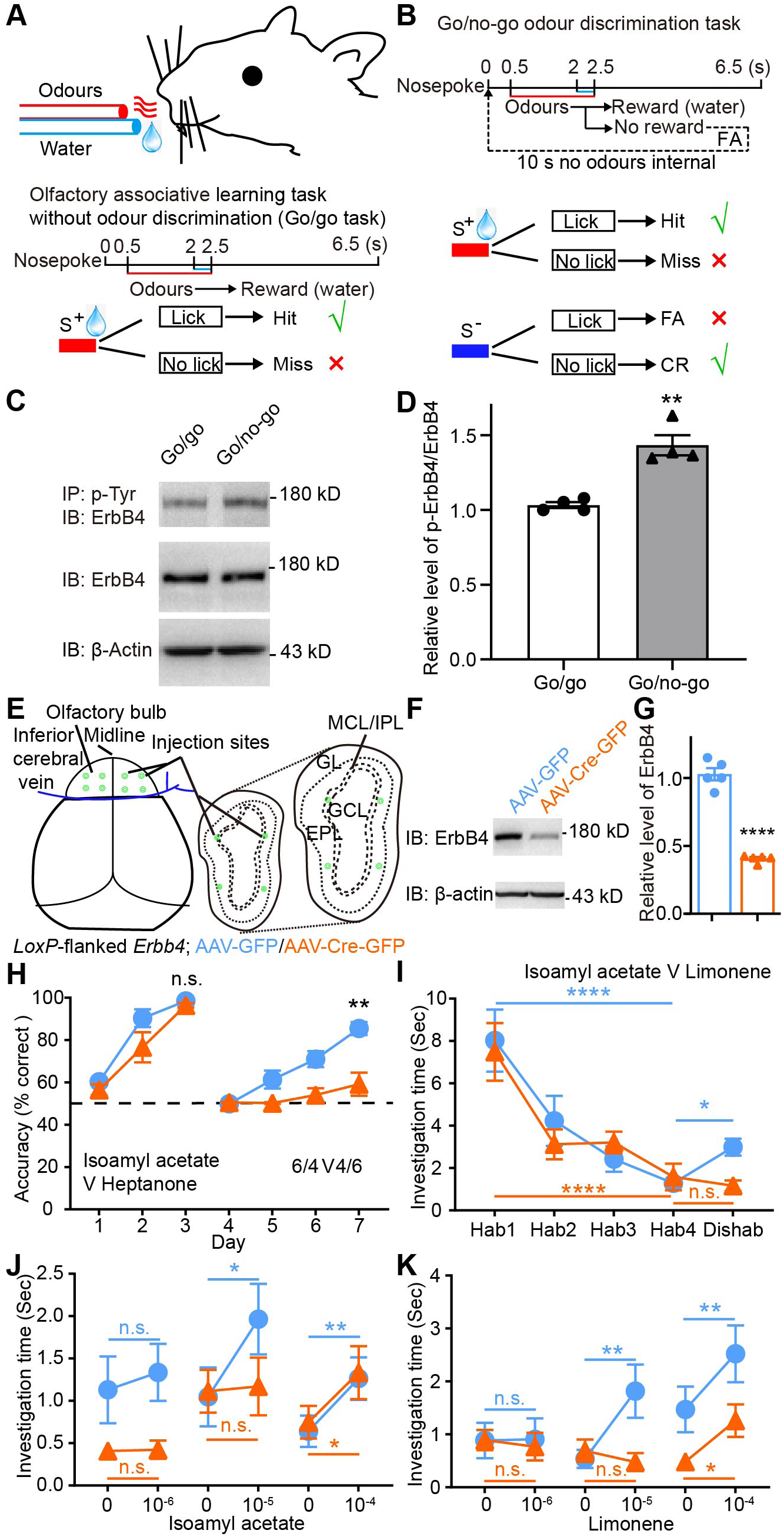
ErbB4 in the OB is critical for odour discrimination and dishabituation. (A) Behavioural paradigm for olfactory associative learning without odour discrimination (go/go task). Mice inserted their snout into the sampling port to trigger odours. The schematic describes the timeline of a single trial. Mice learned to lick the metal tube to receive water in response to either of the odours in the pair (reward, hit). (B) Timeline for a single trial in the go/no-go odour discrimination task. Mice learned to avoid licking the metal tube for the unrewarded odour (correct rejection, CR). Licking when presented with the unrewarded odour (false alarm, FA) led to no water reward and a timeout of up to 10 s. (C and D) ErbB4 activity in the OB was elevated after training on the reinforced go/no-go odour discrimination task. OB cell lysates were subjected to the IP or IB using the indicated antibodies. Relative p-ErbB4/ErbB4 levels were normalized to their respective control groups in the western blot analysis. (*n* = 4 mice per group, *t_(6)_* = 5.75, *P* = 0.0012, unpaired *t* test). (E) Schema indicating virus injection sites. To specifically delete ErbB4 protein in the OB, AAV-Cre-GFP was injected into bilateral OB of neonatal *loxP*-flanked *Erbb4* mice. (F and G) Reduced ErbB4 expression in the OB of a mouse injected with AAV-Cre-GFP (*n* = 5 mice per group, *t_(8)_* = 24.7, *P* < 0.0001, unpaired *t* test). (H) Odour discrimination performance under the reinforced go/no-go task. The accuracy for simple odour pairs was similar for the control and experimental groups (*n* = 8 mice per group, *F_(1, 14)_* = 2.83, *P* = 0.1148, two-way ANOVA). However, the accuracy for difficult odour mixtures (6/4 V 4/6) was reduced in AAV-Cre-GFP mice (*F_(1, 14)_* = 16.14, *P* = 0.0013, two-way ANOVA). (I) Olfactory behavioural performance under a spontaneous habituation/dishabituation task. Both animal groups showed a decline in investigation time to isoamyl acetate over the habituation period (*n* = 9 AAV-GFP mice, *P* < 0.0001; *n* = 10 AAV-Cre-GFP mice, *P* < 0.0001, *F_(3, 51)_* = 21.83, two-way ANOVA). However, AAV-GFP (*F_(1, 17)_* = 3.52, *P* = 0.0065), but not AAV-Cre-GFP mice (*P* = 0.6296, two-way ANOVA), showed an increase in investigation time toward limonene in the dishabituation period. (J) Odour detection threshold to isoamyl acetate. For AAV-GFP mice, the sniffing time toward isoamyl acetate was significantly higher than that for mineral oil at concentrations of 10^-5^ and 10^-4^, but not 10^-6^. (n = 12 mice, *F_(1, 21)_* = 0.19, 4.18 and 14.88, *P* = 0.5689, 0.0111, and 0.0090, two-way ANOVA). These results show that AAV-GFP mice were able to detect isoamyl acetate at a concentration of 10^−5^. For AAV-Cre-GFP mice, the sniffing time toward isoamyl acetate was significantly higher than that for mineral oil at a concentration of 10^-4^, but not at concentrations of 10^-5^ and 10^-6^ (n = 11 mice, *P* = 0.9672, 0.8713, and 0.0172, two-way ANOVA). These results show that AAV-Cre-GFP mice only detect isoamyl acetate at a concentration of 10^−4^. (K) Similarly, AAV-GFP mice were able to detect limonene at a concentration of 10^−5^ (n = 12 mice, *F_(1, 21)_* = 0.03, 4.77 and 12.96, *P* = 0.9603, 0.0011, and 0.0069, two-way ANOVA), whereas AAV-Cre-GFP mice only detected limonene at a concentration of 10^−4^ (n = 11 mice, *P* = 0.7671, 0.5490, and 0.0463, two-way ANOVA). Data are presented as means ± s.e.m. * *P* < 0.05, ** *P* < 0.01, **** *P* < 0.0001, n.s. = not significant. EPL, external plexiform layer; GCL, granule cell layer; GL, glomerular layer; IPL, internal plexiform layer; MCL, mitral cell layer.

**Figure 2.**
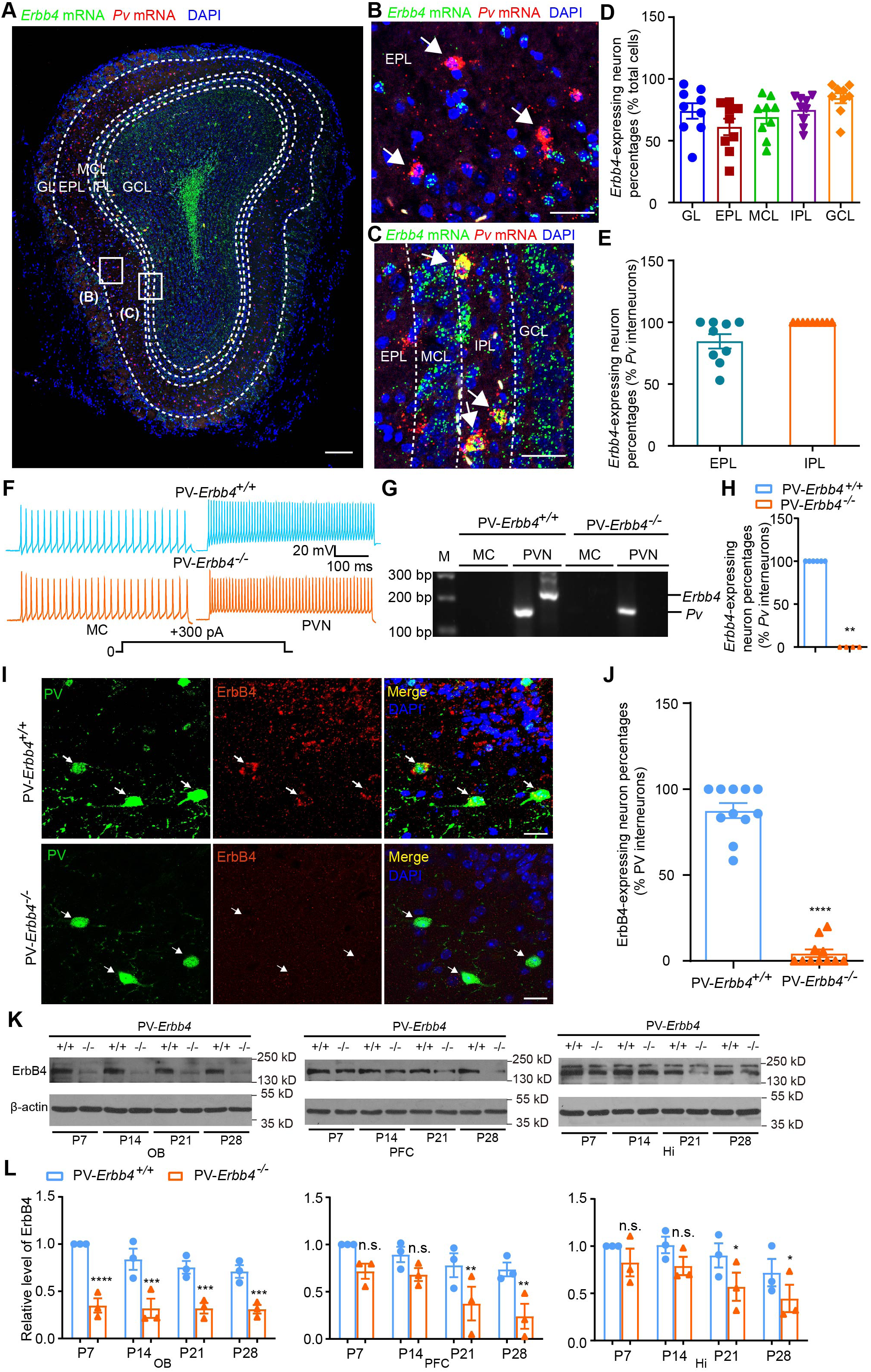
ErbB4 proteins are largely expressed in PV interneurons in the OB. (A) *In vitro* imaging of ErbB4 mRNA in sections from PV-*Erbb4^+/+^* mice. Double single-molecule fluorescence in situ hybridization of ErbB4 (green) and PV (red) in the OB. Scale bar, 200 μm. (B) Magnified view of the EPL box from A. Scale bar, 50 μm. (C) Magnified view of the IPL box from A. Scale bar, 50 μm. (D) Summarized data showing the proportion of ErbB4-expressing neurons in different layers (n = 9 slices from 4 mice). (E) Summarized data showing the proportion of the ErbB4/PV double-positive neurons relative to the total number of PV interneurons in the EPL and IPL (*n* = 9 slices from 4 mice). DAPI staining was used to determine the total number of cells. (F) *In vitro* electrophysiology experiments performed in slices from PV-*Erbb4^+/+^* or PV-*Erbb4^−/−^* mice. Representative examples of action potentials (APs) elicited by positive current injection (500 ms, 300 pA), recorded from an MC (left) and a fast-spiking PV interneuron (right). (G and H) Corresponding single-cell RT-PCR analyses showing that ErbB4 mRNA is detected only in PV interneurons (PVN) from PV-*Erbb4^+/+^* OB (*n* = 6 and 4 neurons from 4 mice, *P* = 0.0048, unpaired *t* test). Dl1000 was used as the size reference (M, 300, 200 and 100 base-pair fragments are indicated). (I and J) Specific deletion of ErbB4 in EPL PV interneurons of the OB (*n* = 11 slices from 4 mice per group, *t_(20)_* = 16.91, *P* < 0.0001, unpaired *t* test). OB sections from PV-*Erbb4^+/+^* and PV-*Erbb4^−/−^* mice (P28) were stained with anti-PV, ErbB4 and DAPI antibody. Scale bars represent 20 μm. (K and L) Western blots showing that ErbB4 in PV-*Erbb4^−/−^* OB was largely reduced from P7 onward, whereas ErbB4 in the PFC and hippocampus began to decrease only at P21. Relative levels were normalized to their respective P7 groups of control littermates (*n* = 3 mice per group, OB: *F_(1, 8)_* = 245.70, *P* < 0.0001, *P* = 0.0002, 0.0006, and 0.0010; PFC: *F_(1, 8)_* = 61.20, *P* = 0.0532, 0.1791, 0.0075, and 0.0021; Hi: *F_(1, 8)_* = 38.25, *P* = 0.2585, 0.1005, 0.0139 and 0.0382, two-way ANOVA). Data are presented as means ± s.e.m. * *P* < 0.05, ** *P* < 0.01, *** *P* < 0.001, **** *P* < 0.001, n.s. = not significant. EPL, external plexiform layer; GCL, granule cell layer; GL, glomerular layer; Hi, hippocampus; MCL, mitral cell layer; PFC, prefrontal cortex.

Therefore, we evaluated whether OB ErbB4 affects the performance of mice in the go/no-go task. Bulbar ErbB4 deficiency was induced by the delivery of adeno-associated virus recombinants expressing Cre recombinase and green fluorescent protein (AAV-Cre-GFP) into both sides of the OB of neonatal mice carrying *loxP*-flanked *Erbb4* alleles (Figure 1E-1G, see Figures 1 and 2 and Supplementary Figure 1C for uncropped western blot images). When learning to discriminate simple odour pairs (Figure 1H), both AAV-GFP– and AAV-Cre-GFP-injected mice reached an accuracy of > 90% within 3 days (∼600 trials), suggesting that ErbB4 in the OB has no influence on learning, memory, or the sense of smell. However, when distinguishing between similar odours, the AAV-GFP group achieved accuracy above 80% after the 4-day training program (∼800 trials), whereas the AAV-Cre-GFP-injected mice showed significantly lower accuracy. This indicates that ErbB4 in the OB maintains the ability of mice to discriminate between complex odours.

We conducted a habituation/dishabituation test to assess the spontaneous behavioural responses of mice to repeatedly presented odours in the absence of any reward (Figure 1I) to further determine the role of the OB ErbB4 in olfactory behaviours. Both AAV-GFP and AAV-Cre-GFP groups were habituated to four successive odour presentations with 2-min intertrial intervals, and their behavioural responses to the same odour (isoamyl acetate) were progressively reduced. AAV-GFP mice rapidly dishabituated, and their investigation time increased when a novel odour was presented (limonene, fifth trial), whereas AAV-Cre-GFP mice did not show significantly altered behavioural responses to the novel odour. Additionally, the olfactory detection thresholds for isoamyl acetate and limonene were approximately 10^-5^ in AAV-GFP mice, but 10^-4^ in AAV-Cre-GFP mice (Figure 1J and 1K). In the habituation and dishabituation tasks, the two odours were used at levels far above the threshold.

These results demonstrate that ErbB4 in the OB contributes to olfactory discrimination and dishabituation but not to olfactory memory and habituation.

### ErbB4 is expressed in PV interneurons within the EPL and IPL layers of the OB

RNAscope analysis was performed to detect mRNA levels of *Erbb4* in the OB of PV-*Erbb4^+/+^* mice. As shown in Figure 2A-2D, *Erbb4* mRNA was present in all five layers of the OB in PV-*Erbb4^+/+^*mice (green); *Erbb4*-expressing neurones accounted for 74.0%, 61.2%, 69.1%, 74.9%, and 84.4% of the total cells (DAPI, blue) in the glomerular layer (GL), EPL, mitral cell layer (MCL), IPL, and granule cell layer (GCL), respectively (Figure 2D). PV interneurons (red) in the OB were primarily localised to the EPL and IPL (Figure 2A), where *Erbb4* mRNA tended to be located within PV interneurons (yellow) (Figure 2A-2C). Of the PV interneurons in the EPL and IPL, 84.6% and 100% expressed *Erbb4*, respectively (Figure 2E). Overall, *Erbb4* mRNA levels in PV interneurons were the highest in the IPL and lowest in the EPL.

To further ascertain *Erbb4* mRNA expression of individual PV interneurons within EPL and IPL, we conducted patch-clamp electrophysiology followed by single-cell analysis of reverse transcription-polymerase chain reaction (RT-PCR) in OB slices from PV-*Erbb4^+/+^*and PV-ErbB4 conditional knockout (PV-*Erbb4^−/−^*) mice. In addition to their morphology, size, and location, MCs were electrophysiologically characterised as having low-frequency spike discharges (<100 Hz). In contrast, PV interneurons were identified based on their high-frequency spike discharges (>100 Hz) (Figure 2F). The cytosolic contents from identified MCs and PV neurones were then subjected to RT-PCR, showing that *Erbb4* mRNA was only present in PV interneurons from PV-*Erbb4^+/+^*OB, but not in any of the MCs or PV interneurons from PV-*Erbb4^−/−^*OB (Figure 2G and 2H).

Immunofluorescence analysis showed that ErbB4 immunoreactivity within the EPL was present in PV interneurons of PV-*Erbb4^+/+^* mice; this was largely diminished in PV-*Erbb4^−/−^* mice (Figure 2I and J). Western blots showed that the levels of ErbB4 proteins were largely reduced, but not abolished, in the OB, prefrontal cortex (PFC), and hippocampus (Hi) of PV-*Erbb4^−/−^* mice compared with their control littermates (Figure 2K and 2L, see Figures 1 and 2 and Supplementary Figure 1D for uncropped western blot images). This suggests that ErbB4 is largely expressed in the OB PV interneurons. Interestingly, ErbB4 deletion in the OB occurred at least 2 weeks prior to deletion in the prefrontal cortex and hippocampus (Figure 2K and 2L). Nissl staining showed that the overall lamina structures of the OB were not affected in PV-*Erbb4^−/−^* mice (Figure 2 and Supplementary Figure 2).

These results demonstrate the strong expression of ErbB4 in PV fast-spiking interneurons within the EPL and IPL layers of the OB.

### ErbB4 in PV interneurons is necessary for olfactory behaviours

To determine the contribution of ErbB4 expressed in PV interneurons to olfactory behaviours, we conducted reinforced go/no-go tests of olfactory discrimination in PV-*Erbb4^+/+^*and PV-*Erbb4^−/−^* mice. Both groups learned to distinguish distinct odour pairs with more than 90% accuracy within 600 trials (Figure 3A); PV-*Erbb4^−/−^*mice exhibited significantly lower accuracy than their control littermates for similar odour pairs (Figure 3A). During habituation/dishabituation tasks, both groups of mice habituated to the successive presentation of a familiar odour (isoamyl acetate, fourth trial) (Figure 3B); when a novel odour (limonene, fifth trial) was presented, PV-*Erbb4^−/−^* mice did not increase their investigation time, unlike their control littermates (Figure 3B). Similarly, when the order of odour presentation was reversed, the PV-*Erbb4^−/−^* group showed impaired dishabituation, whereas the PV-*Erbb4^+/+^* group did not (Figure 3C). The olfactory detection thresholds for isoamyl acetate and limonene were slightly higher in PV-*Erbb4^−/−^* mice than in PV-*Erbb4^+/+^* mice (Figure 3D and 3E).

**Figure 3.**
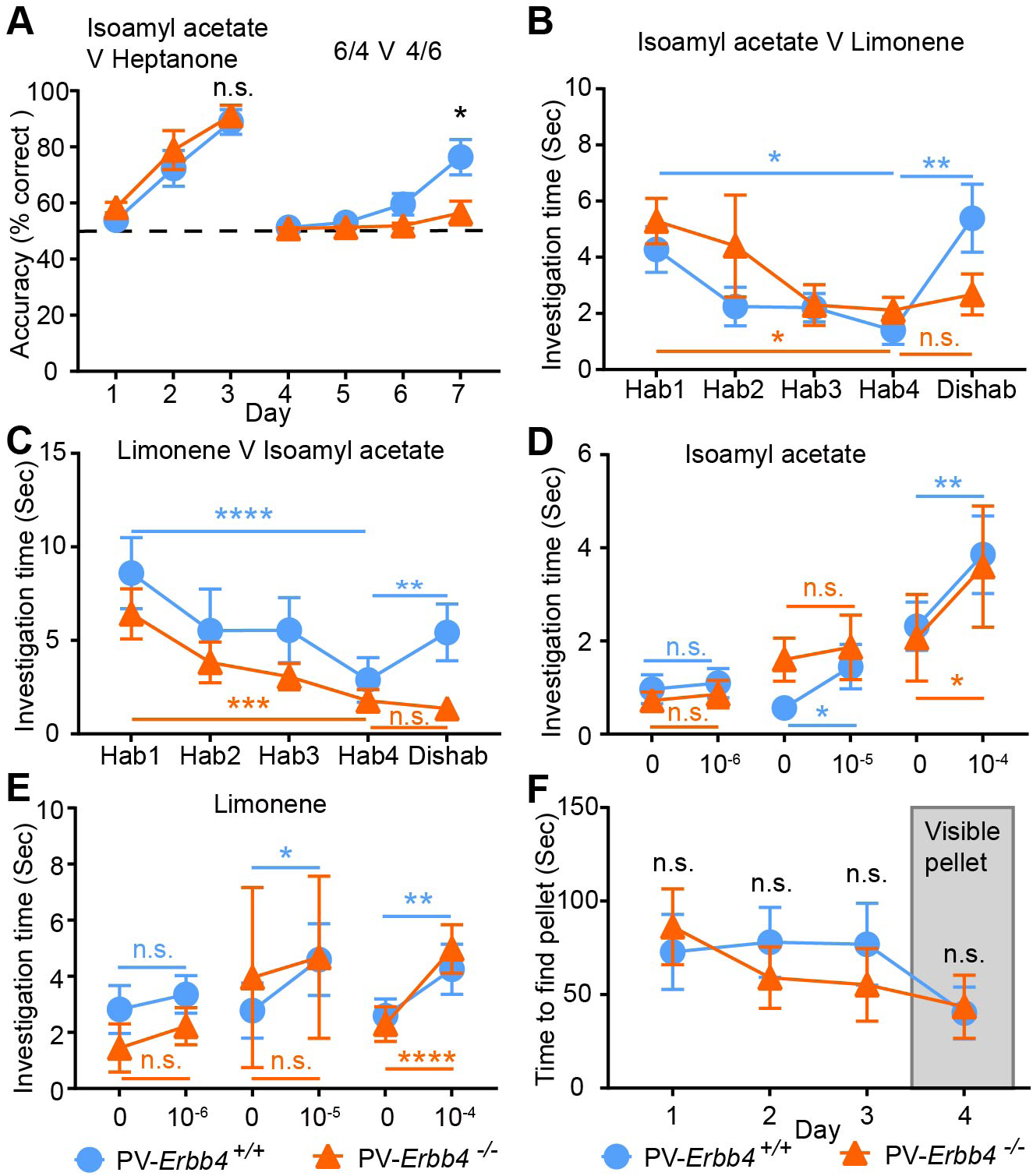
ErbB4 in PV interneurons is critical for olfactory behaviours. (A) Odour discrimination under the reinforced go/no-go task in PV-*Erbb4^−/−^* mice. The accuracy for simple odour pairs was indistinguishable (*n* = 5 and 6 mice, *F_(1, 9)_* = 0.70, *P* = 0.4260, two-way ANOVA). However, the accuracy for difficult odour pairs was significantly lower in PV-*Erbb4^−/−^*mice (*F_(1, 9)_* = 9.12*, P* = 0.0144, two-way ANOVA). (B) Olfactory behavioural performance under the spontaneous habituation/dishabituation task. Both animal groups habituated to isoamyl acetate (*n* = 12 mice per group, *F_(3, 66)_* = 6.68, *P* = 0.0349 and 0.0164, two-way ANOVA). However, PV-*Erbb4^+/+^*mice (*F_(1, 22)_* = 8.93, *P* = 0.0025), but not PV-*Erbb4^−/−^*mice (*P* = 0.8451, two-way ANOVA), dishabituated to limonene. (C) Olfactory behavioral performance under the reversed habituation/dishabituation task. Both animal groups habituated to limonene (*F_(3, 54)_* = 16.33, *P* < 0.0001 and 0.0003, two-way ANOVA). However, PV-*Erbb4^+/+^* mice (*F_(1, 18)_* = 5.22, *P* = 0.0023), but not PV-*Erbb4^−/−^*mice (*P* = 0.7890, two-way ANOVA), dishabituated to isoamyl acetate. (D) PV-*Erbb4^+/+^*mice were able to detect isoamyl acetate at a concentration of 10^−5^ (*n* = 10 mice, *F_(1, 18)_* = 0.60, 3.74 and 16.69, *P* = 0.6069, 0.0498, and 0.0096 for 10^-6^, 10^-5^, and 10^-4^, two-way ANOVA). PV-*Erbb4^−/−^* mice only detected isoamyl acetate at a concentration of 10^−4^ (*n* = 10 mice, *P* = 0.5764, 0.5353, and 0.0100 for 10^-6^, 10^-5^, and 10^-4^, two-way ANOVA). (E) PV-*Erbb4^+/+^* mice could detect limonene at a concentration of 10^−5^ (*n* = 10 mice, *F_(1, 18)_* = 1.75, 7.95 and 32.51, *P* = 0.4540, 0.0106, and 0.0064 for 10^-6^, 10^-5^, and 10^-4^, two-way ANOVA). PV-*Erbb4^−/−^* mice only detected limonene at a concentration of 10^−4^ (*n* = 10 mice, *P* = 0.2842, 0.2709, and < 0.0001 for 10^-6^, 10^-5^, and 10^-4^, two-way ANOVA). (F) The latency for mice to locate the buried and visible food pellets did not differ between the groups (*n* = 14 and 11 mice, for the buried food pellet, *F_(1, 23)_* = 0.21, *P* = 0.9786, 0.9279, and 0.8873; for the visible food pellet, *P* > 0.9999, two-way ANOVA). Data are presented as means ± s.e.m. * *P* < 0.05, ** *P* < 0.01, *** *P* < 0.001, **** *P* < 0.0001, n.s. = not significant.

We also tested mice using the buried food test, which depends on the animal’s natural tendency to utilise olfactory cues for foraging, to confirm whether the mice could smell volatile odours. As shown in Figure 3F, both PV-*Erbb4^+/+^* and PV-*Erbb4^−/−^* mice found the buried pellets rapidly over three consecutive testing days. On trial day 4, the pellet was easily located within a similar timeframe as when the food pellet was made visible by placing it on the surface. This suggested that ErbB4 in PV interneurons is not required for the ability of mice to detect and locate odorous sources.

These findings reveal that ErbB4 signalling in PV interneurons serves as an essential regulator of olfactory discrimination and dishabituation.

### ErbB4 in PV interneurons mediates odour-evoked responses of M/TCs in the OB

To investigate the overall functional relevance of ErbB4 signalling in PV interneurons in regulating M/TC responses, we recorded extracellular spontaneous and odour-evoked activity from M/TCs in the OB via tetrodes implanted in awake, head-fixed PV-*Erbb4^−/−^* and PV-*Erbb4*^+/+^ mice (Figure 4A). To obtain individual neuronal spikes, units were identified using principal component analysis (PCA) scan clustering of the spikes (Figure 4 and Supplemental Figure 1A and 1B). ErbB4 deletion in PV interneurons did not directly affect the spike waveform of M/TCs *in vivo*, with no significant differences in peak-to-valley ratio or half-valley width between PV-*Erbb4^−/−^* and PV-*Erbb4^+/+^*mice (Figure 4, Supplementary Figure 1C–1E). Both excitatory (upper, odour-evoked firing rate > spontaneous firing rate) and inhibitory (lower, odour-evoked firing rate < spontaneous firing rate) responses were observed upon odour presentation using the odour delivery system (Figure 4B and 4C). Figure 4D shows the heat maps of odour-evoked changes in mean firing rate (ΔMFR) of 240 M/TCs in PV-*Erbb4^+/+^* mice (left) and 232 M/TCs in PV-*Erbb4^−/−^* mice (right). As shown in Figure 4E, spontaneous firing of M/TCs was similar in PV-*Erbb4^+/+^* and PV-*Erbb4^−/−^* groups; odour-evoked firing of M/TCs was reduced by ErbB4 deletion in PV interneurons. Both the absolute value of odour-evoked changes in mean firing rate (ΔMFR) and the signal-to-noise ratio (SNR) were lower in PV-*Erbb4^−/−^* M/TCs than in PV-*Erbb4^+/+^* M/TCs (Figure 4E).

**Figure 4.**
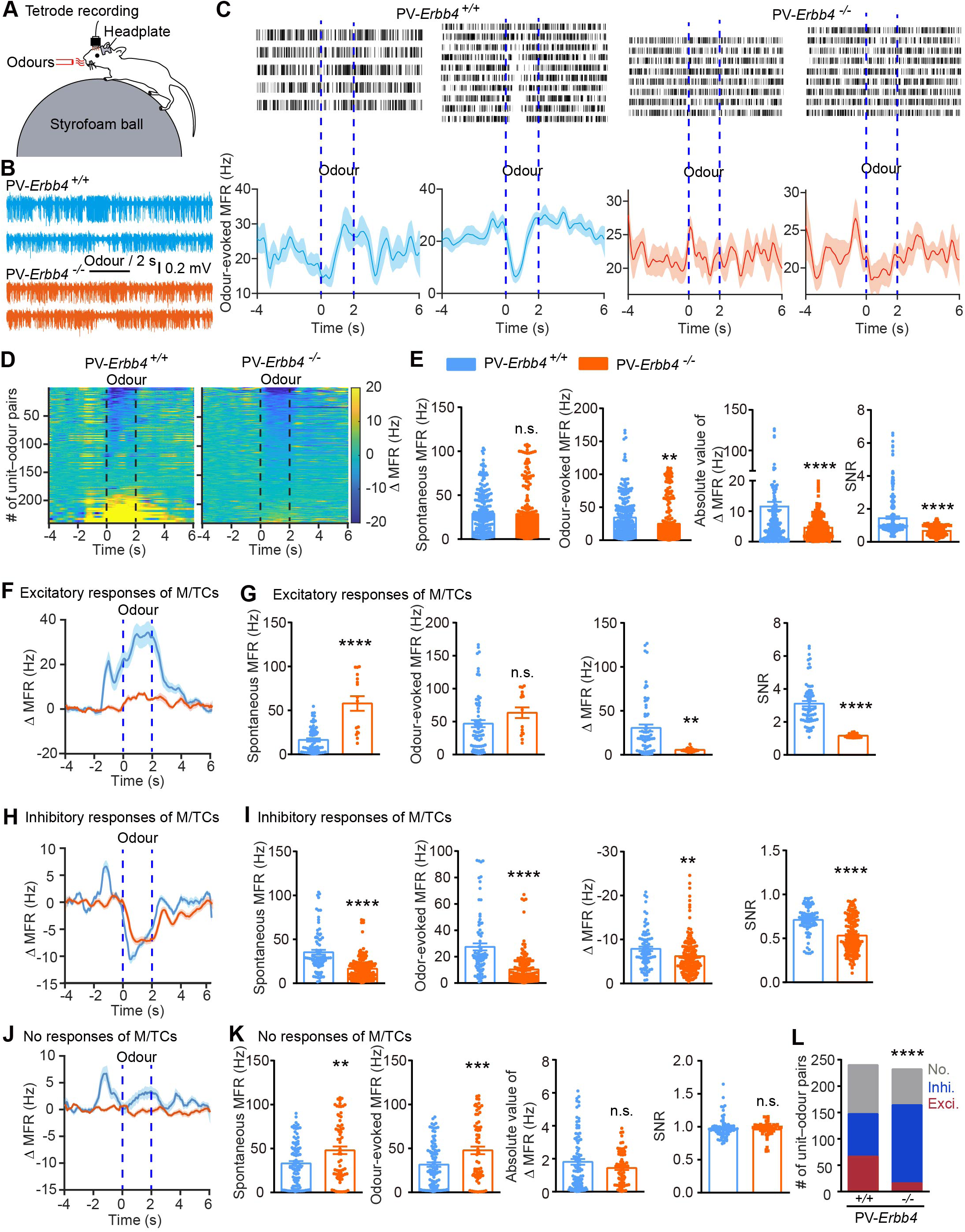
Odour-evoked responses in M/TCs are decreased in PV-*Erbb4^−/−^* mice. (A) Schematic of *in vivo* odour-evoked electrophysiological recordings in awake, head-fixed mice with ErbB4 knocked out in PV interneurons. (B) Representative raw traces of spike activity before (spontaneous), during (odour-evoked), and after 2-s odour stimulation in PV-*Erbb4^+/+^* and PV-*Erbb4^−/−^* mice. (C) Examples of raster plots (top) and PSTHs of the firing rate (bottom) for odour-evoked excitatory (left) and inhibitory (right) responses in PV-*Erbb4^+/+^*and PV-*Erbb4^−/−^* mice. PSTHs were smoothed with a Gaussian filter with a standard deviation of 1500 ms. (D) Heat maps of the odour-evoked changes in mean firing rate (ΔMFR) across all unit–odour pairs in PV-*Erbb4^+/+^*mice (*n* = 240 unit–odour pairs from 4 mice) and PV-*Erbb4^−/−^*mice (*n* = 232 unit–odour pairs from 4 mice). (E) Quantitative analysis of the spontaneous firing rate (*t_(470)_* = 0.27, *P* = 0.7899), odour-evoked MFR (*t_(470)_* = 3.30, *P* = 0.0010), absolute value of odour-evoked changes (*t_(470)_* = 5.05, *P* < 0.0001), and SNR (*t_(470)_* = 9.29, *P* < 0.0001, unpaired *t* test) across all unit–odour pairs. (F) Odour-evoked excitatory changes in MFR (ΔMFR) for M/TCs recorded from PV-*Erbb4^+/+^* mice (*n* = 66 unit–odour pairs from 4 mice) and PV-*Erbb4^−/−^* mice (*n* = 16 unit–odour pairs from 4 mice). (G) Quantitative analysis of the spontaneous firing rate (*t_(80)_* = 7.67, *P* < 0.0001), odour-evoked MFR (*t_(80)_* = 1.43, *P* = 0.1578), odour-evoked ΔMFR (*t_(80)_* = 3.10, *P* = 0.0027), and SNR (*t_(80)_* = 5.91, *P* < 0.0001, unpaired *t* test) across excitatory unit–odour pairs. (H) Odour-evoked inhibitory ΔMFR for M/TCs recorded from PV-*Erbb4^+/+^*mice (*n* = 81 unit–odour pairs from 4 mice) and PV-*Erbb4^−/−^*mice (*n* = 148 unit–odour pairs from 4 mice). (I) Quantitative analysis of spontaneous firing rate (*t_(227)_* = 7.52, *P* < 0.0001), odour-evoked MFR (*t_(227)_* = 7.23, *P* < 0.0001), odour-evoked ΔMFR (*t_(227)_* = 2.72, *P* = 0.0070), and SNR (*t_(227)_* = 6.91, *P* < 0.0001, unpaired *t* test) across inhibitory unit–odour pairs. (J) ΔMFR for units with no response to odour in PV-*Erbb4^+/+^* mice (*n* = 93 unit–odour pairs from 4 mice) and PV-*Erbb4^−/−^* mice (*n* = 68 unit–odour pairs from 4 mice). (K) Quantitative analysis of spontaneous firing rate (*t_(159)_* = 3.13, *P* = 0.0021), odour-evoked MFR (*t_(159)_* = 3.38, *P* = 0.0009), absolute value of ΔMFR (*t_(159)_* = 1.63, *P* = 0.1057), and SNR (*t_(159)_* = 0.43, *P* = 0.6687, unpaired *t* test) across “no response” unit–odour pairs. (L) Distribution of excitatory, inhibitory, and “no response” units in PV-*Erbb4^+/+^* and PV-*Erbb4^−/−^* mice (*χ^2^_(2)_* =53.85, *P* < 0.0001, Chi-Square tests). ** *P* < 0.01, *** *P* < 0.001, **** *P* < 0.0001, n.s. = not significant.

We next analysed the differences in M/TCs with excitatory responses (Figure 4F and 4G), inhibitory responses (Figure 4H and 4I), and no responses (Figure 4J and 4K) to odour presentation in PV-*Erbb4^+/+^* and PV-*Erbb4^−/−^* mice. Compared with the PV-*Erbb4^+/+^* group, PV-*Erbb4^−/−^* M/TCs had significantly higher spontaneous firing rates for excitatory responses (Figure 4G). The odour-evoked firing rate was not changed by ErbB4 deletion in PV interneurons. The odour-evoked changes in ΔMFR and SNR markedly decreased in PV-*Erbb4^−/−^* M/TCs (Figure 4F and 4G). Compared with the PV-*Erbb4^+/+^*group, M/TCs in PV-*Erbb4^−/−^* mice exhibited significantly lower spontaneous and odour-evoked inhibitory firing rates (Figure 4I). Interestingly, the odour-evoked changes in ΔMFR and SNR also showed a slight decrement in PV-*Erbb4^−/−^* M/TCs (Figure 4H and 4I). Spontaneous and odour-evoked firing rates for M/TCs classes that had no responses during odour presentation were higher in PV-*Erbb4^−/−^* group than in the PV-*Erbb4^+/+^* group (Figure 4K). The absolute values of odour-evoked changes (ΔMFR) and SNR showed no differences between the two groups (Figure 4J and 4K). Furthermore, the distribution of types of odour-evoked responses was significantly altered in PV-*Erbb4*^−/−^ mice compared with PV-*Erbb4*^+/+^ mice (Figure 4L), suggesting compromised odour detection.

These results demonstrate that ErbB4 signalling in PV interneurons mediates the fine-tuning of M/TC odour-evoked responses.

### ErbB4 in PV interneurons decreases the power of the ongoing oscillations

Next, we investigated the overall functional relevance of ErbB4 in PV interneurons by examining whether their deletion affects spontaneous oscillations in the OB. Raw local field potential (LFP) signals were divided into different frequency bands: theta (2–12 Hz), beta (15–35 Hz), low gamma (36–65 Hz), and high gamma (66–95 Hz) (Figure 5A). For all frequency bands of the ongoing baseline LFP, the power was higher in PV-*Erbb4^−/−^* mice than in PV-*Erbb4^+/+^* mice (Figure 5B-5I), suggesting that ErbB4 signalling in PV interneurons controls the spontaneous ongoing network oscillations in the OB.

**Figure 5.**
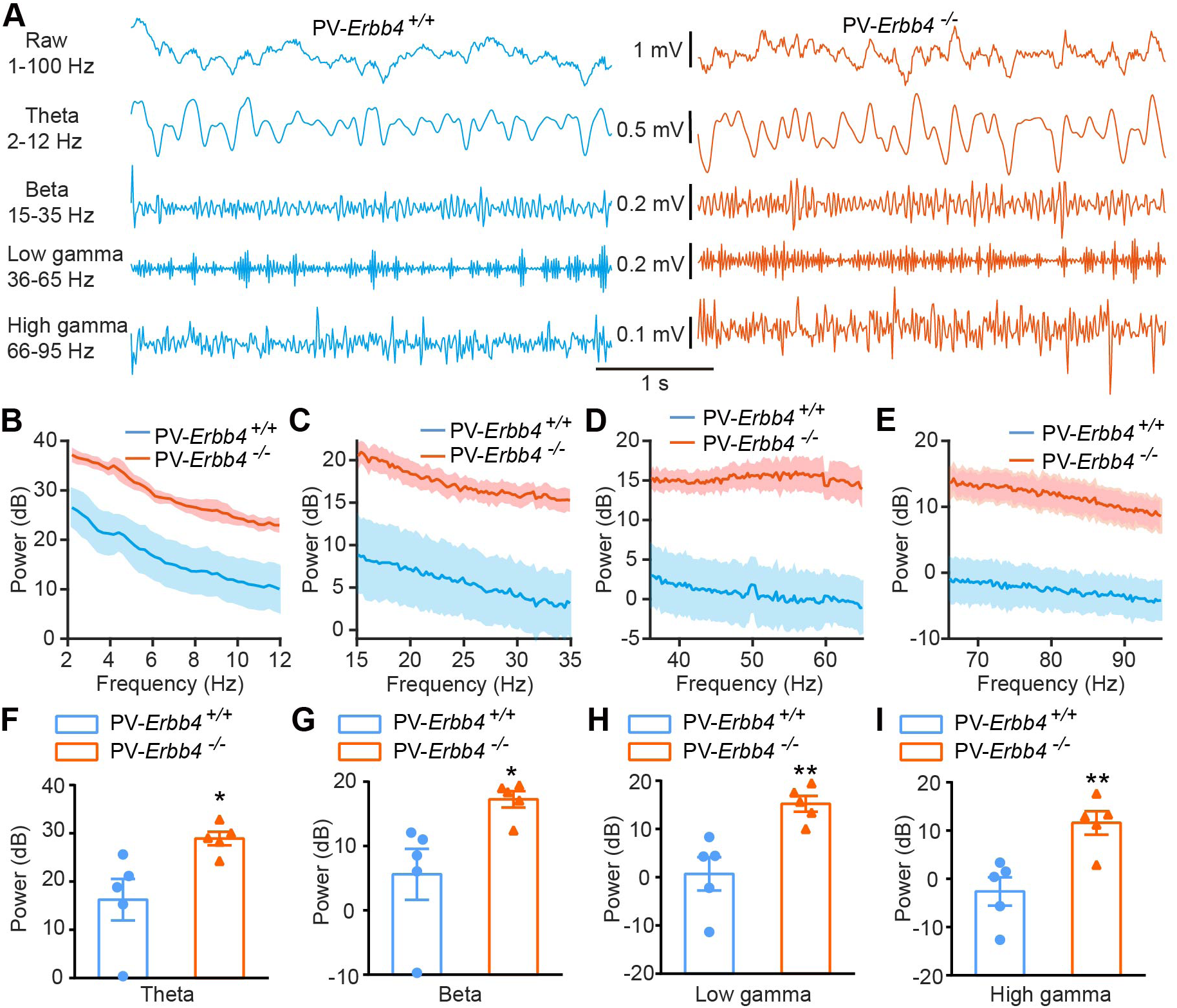
The ongoing LFP in the OB is increased in PV-*Erbb4^−/−^* mice. (A) Examples of ongoing LFP signals recorded in the OB from PV-*Erbb4^+/+^*and PV-*Erbb4^−/−^* mice. The five rows show the raw traces and the filtered theta, beta, low-gamma, and high-gamma signals. (B-E) Quantitative analysis of the averaged power spectra in the theta, beta, low-gamma, and high-gamma bands for the two groups. (F-I) Comparisons of power in the theta (*n* = 5 mice per group, *t_(8)_* = 2.80, *P* = 0.0233, unpaired *t* test), beta (*t_(8)_* = 2.80, *P* = 0.0232, unpaired *t* test), low-gamma (*t_(8)_* = 3.80, *P* = 0.0053, unpaired *t* test), and high-gamma (*t_(8)_* = 3.73, *P* = 0.0058, unpaired *t* test) oscillations in the OB in the two groups. * *P* < 0.05, ** *P* < 0.01.

### ErbB4 in PV interneurons regulates the OB output

To further evaluate the physiological role of the PV interneuron ErbB4 in the regulation of OB output, we recorded the spontaneous and evoked excitatory firing rates from the principal MCs (output cells) in OB slices. Cell-attached recordings showed that the frequency of MC spontaneous action potentials (sAPs) was dramatically enhanced in PV-*Erbb4^−/−^* mice compared with PV-*Erbb4^+/+^* mice (Figure 6A and 6B). Furthermore, the frequency of evoked action potentials (eAPs) was also higher in PV-*Erbb4^−/−^* mice (Figure 6A and 6B). However, the ratio of eAP to sAP firing (i.e., the SNR) was significantly lower in PV-*Erbb4^−/−^* mice (Figure 6A and 6B). More importantly, the SNR in response to stimuli at various intensities was lower in PV-*Erbb4^−/−^* mice than in their control littermates (Figure 6C and 6D), even though the overall current-evoked activity was elevated in PV-*Erbb4^−/−^* mice compared with their control littermates (Figure 6E and 6F).

**Figure 6.**
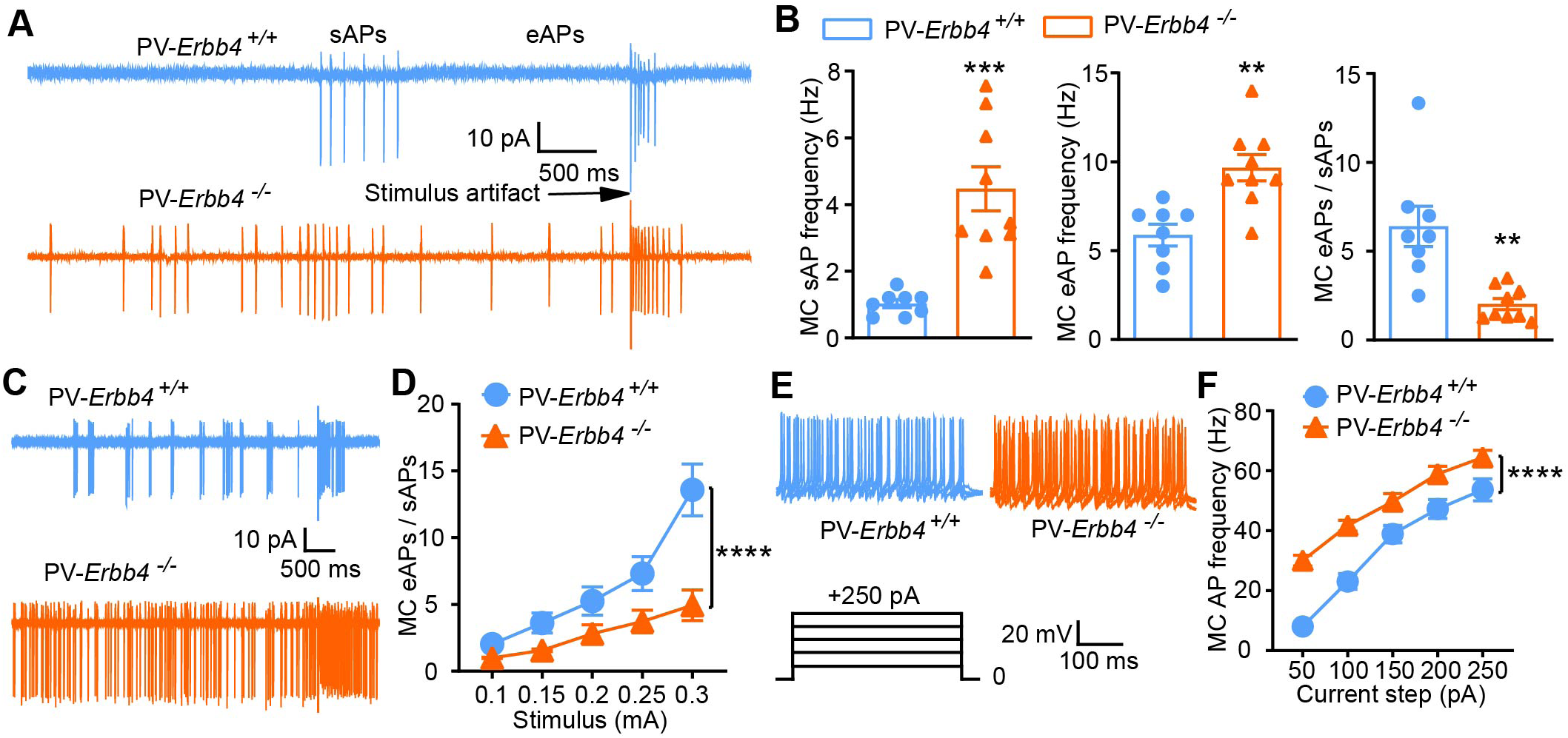
PV-*Erbb4^−/−^* mice have impaired information output from the OB. (A and B) Increased frequency of MC sAPs and olfactory nerve-evoked APs (eAPs) in PV-*Erbb4^−/−^* mice, but decreased ratio of eAPs to sAPs (*n* = 8-9 from 3 mice per group, sAPs: *t_(15)_* = 4.84, *P* = 0.0002, eAPs: *t_(15)_* = 3.87, *P* = 0.0015; ratio: *t_(15)_* = 3.90, *P* = 0.0014, unpaired *t* test). (C and D) The eAP-to-sAP ratio was reduced in PV-*Erbb4^−/−^* mice as the intensity of the stimulus increased (*n* = 8 from 3 mice per group, *F_(1, 70)_* = 32.39, *P* < 0.0001, two-way ANOVA). (E and F) AP frequency elicited by injection of positive currents was higher in PV-*Erbb4^−/−^* mice than PV-*Erbb4^+/+^* mice (*n* = 9 from 4 and 3 mice per group, *F_(1, 80)_* = 78.9, *P* < 0.0001, two-way ANOVA). Data are presented as means ± s.e.m. ** *P* < 0.01, *** *P* < 0.001, **** *P* < 0.0001.

These data indicate an involvement of PV interneuron ErbB4 in OB odour information processing.

### ErbB4 in PV interneurons maintains inhibitory circuit activity in the OB

To investigate the neural mechanisms underlying the observed hyperexcitability in PV-*Erbb4^−/−^* mouse MCs, we tested whether the effect of ErbB4 deletion in PV interneurons could be occluded by GABAergic blockage. Bicuculline (10 μM), a selective GABA_A_ receptor antagonist, elevated sAP firing in MCs from PV-*Erbb4^+/+^* mice but did not further increase MC hyperexcitability in PV-*Erbb4^−/−^* mice (Figure 7A and 7B). Bicuculline also reduced the eAP:sAP ratio in PV-*Erbb4^+/+^* MCs but not in PV-*Erbb4^−/−^*MCs (Figure 7A and 7B), suggesting that the impaired output of PV-*Erbb4^−/−^* MCs depends on GABAergic downregulation. Therefore, we recorded GABA_A_ receptor-mediated miniature inhibitory postsynaptic currents (mIPSCs) from MCs in whole-cell mode. As shown in Figure 7C and 7D, the mIPSC frequency, but not the amplitude, was significantly reduced in PV-*Erbb4^−/−^* MCs, suggesting a presynaptic impairment of inhibitory transmission. Neither the frequency nor the amplitude of glutamatergic receptor-mediated miniature excitatory postsynaptic currents (mEPSCs) was altered in PV-*Erbb4^−/−^*MCs compared with their control littermates (Figure 7E and 7F). In contrast, both the frequency and amplitude of mEPSCs in OB PV interneurons were markedly reduced in PV*-Erbb4^−/−^* mice (Figure 7, Supplementary Figure 1A and 1B). Further analyses of PV interneuron intrinsic properties showed increased action potential frequency in PV-*Erbb4^⁻/^*^⁻^ mice. In contrast, action potential amplitude and half-width remained unaltered (Figure 7, Supplementary Figure 1C-1E), indicating preserved spike waveform following ErbB4 deletion. These results suggest that ErbB4 signalling in PV interneurons regulates olfactory information output by maintaining GABAergic inputs to MCs in the OB.

**Figure 7.**
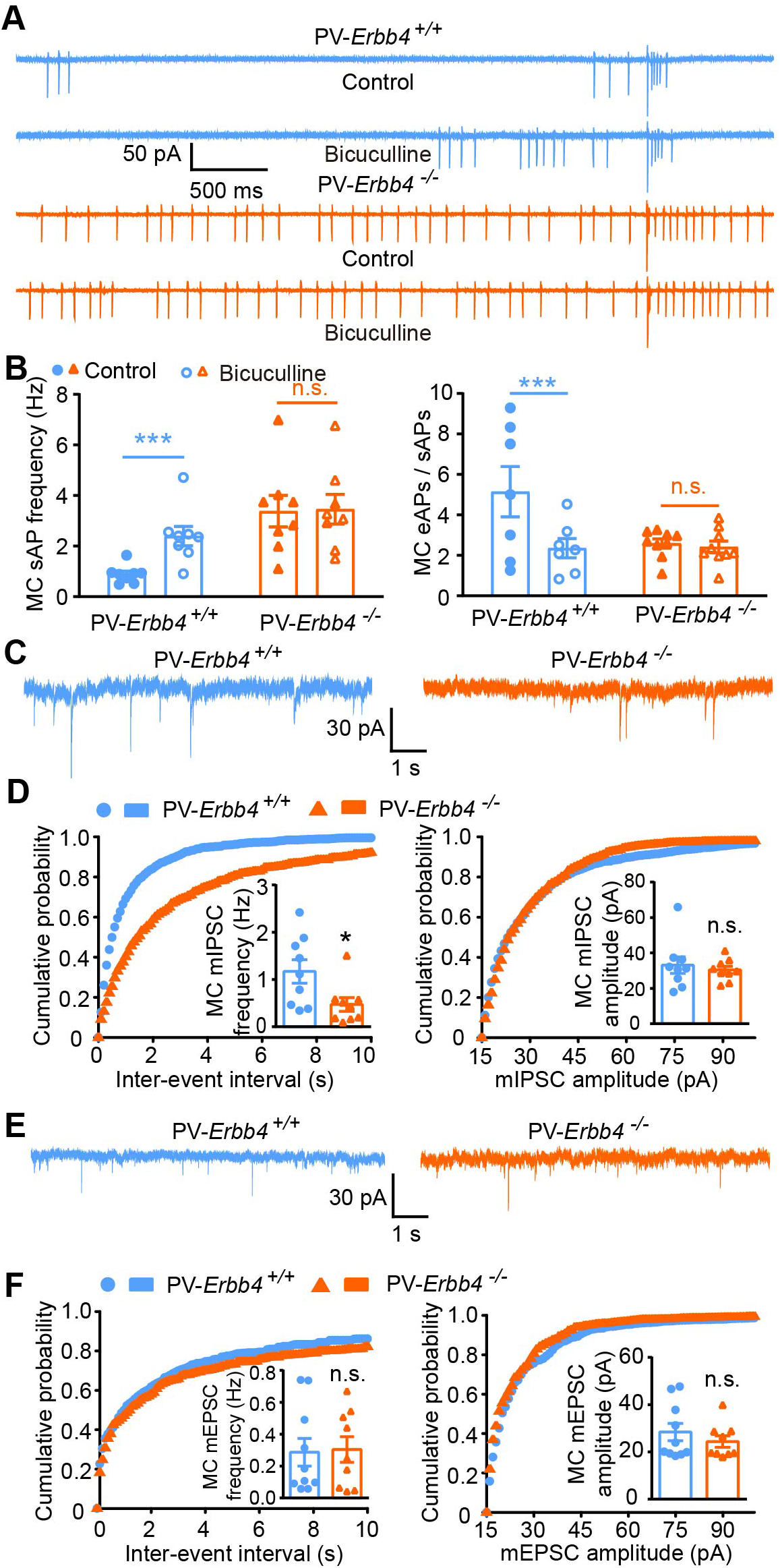
Decreased GABAergic transmission mediates the OB output impairments in PV-*Erbb4^−/−^* mice. (A and B) The increased sAP frequency and decreased ratio of eAPs to sAPs in MCs could not be further enhanced by bicuculline in PV-*Erbb4^−/−^* mice (*n* = 7 from 3 PV-*Erbb4^+/+^* mice, *n* = 9 from 3 PV-*Erbb4^−/−^* mice; for sAP, *F_(1, 14)_* = 17.08, *P* = 0.0008 and 0.5531; for ratio of eAPs to sAPs, *F_(1, 14)_* = 13.99, *P* = 0.0007 and 0.9360, two-way ANOVA). (C and D) The frequency but not the amplitude of MC mIPSCs was lower in PV-*Erbb4^−/−^* mice (*n* = 9 from 7 PV-*Erbb4^+/+^*and 6 PV-*Erbb4^−/−^*mice; for frequency, *t_(16)_* = 2.45, *P* = 0.0263; for amplitude, *t_(16)_* = 0.53, *P* = 0.6038, unpaired *t* test). (E and F) Neither frequency nor amplitude of MC mEPSCs was different in PV-*Erbb4^−/−^* mice versus PV-*Erbb4^+/+^*mice (*n* = 10 from 5 PV-*Erbb4^+/+^* mice, *n* = 9 from 7 PV-*Erbb4^−/−^* mice; for frequency, *t_(17)_* = 0.14, *P* = 0.8899; for amplitude, *t_(17)_* = 0.89, *P* = 0.3863, unpaired *t* test). Data are presented as means ± s.e.m. * *P* < 0.05, *** *P* < 0.001, **** *P* < 0.0001, n.s. = not significant.

Next, we examined the effect of a specific PV-ErbB4 knockout on the recurrent inhibition of MCs. Prolonged hyperpolarisation after AP firing, which resulted from recurrent inhibition, was observed in the whole-cell patch-clamp configuration. Although the strength of recurrent inhibition is reported to depend on MC firing rates (Margrie et al., 2001), neither the amplitude nor the decay times of recurrent inhibitory postsynaptic potentials (IPSPs) showed significant alteration with the increment in AP frequency in PV-*Erbb4^−/−^* mice (Figure 8A and 8B). One possible explanation is that the loss of ErbB4 in the PV interneurons severely impedes recurrent IPSPs. To test this hypothesis, we recorded recurrent IPSPs after a given frequency of APs in PV-*Erbb4^−/−^* mice. As shown in Figure 8C and Figure 8 and Supplementary Figure 1A, a current injection of 80 pA in PV-*Erbb4^−/−^* mice elicited approximately the same firing rate as a current injection of 100 pA in PV-*Erbb4^+/+^* mice. Remarkably, both the amplitude and the decay time constant of the subsequent recurrent IPSP were reduced in PV-*Erbb4^−/−^*mice (Figure 8D). Thus, ErbB4-expressing PV interneurons mediate recurrent inhibition induced by the same MC firing intensity.

**Figure 8.**
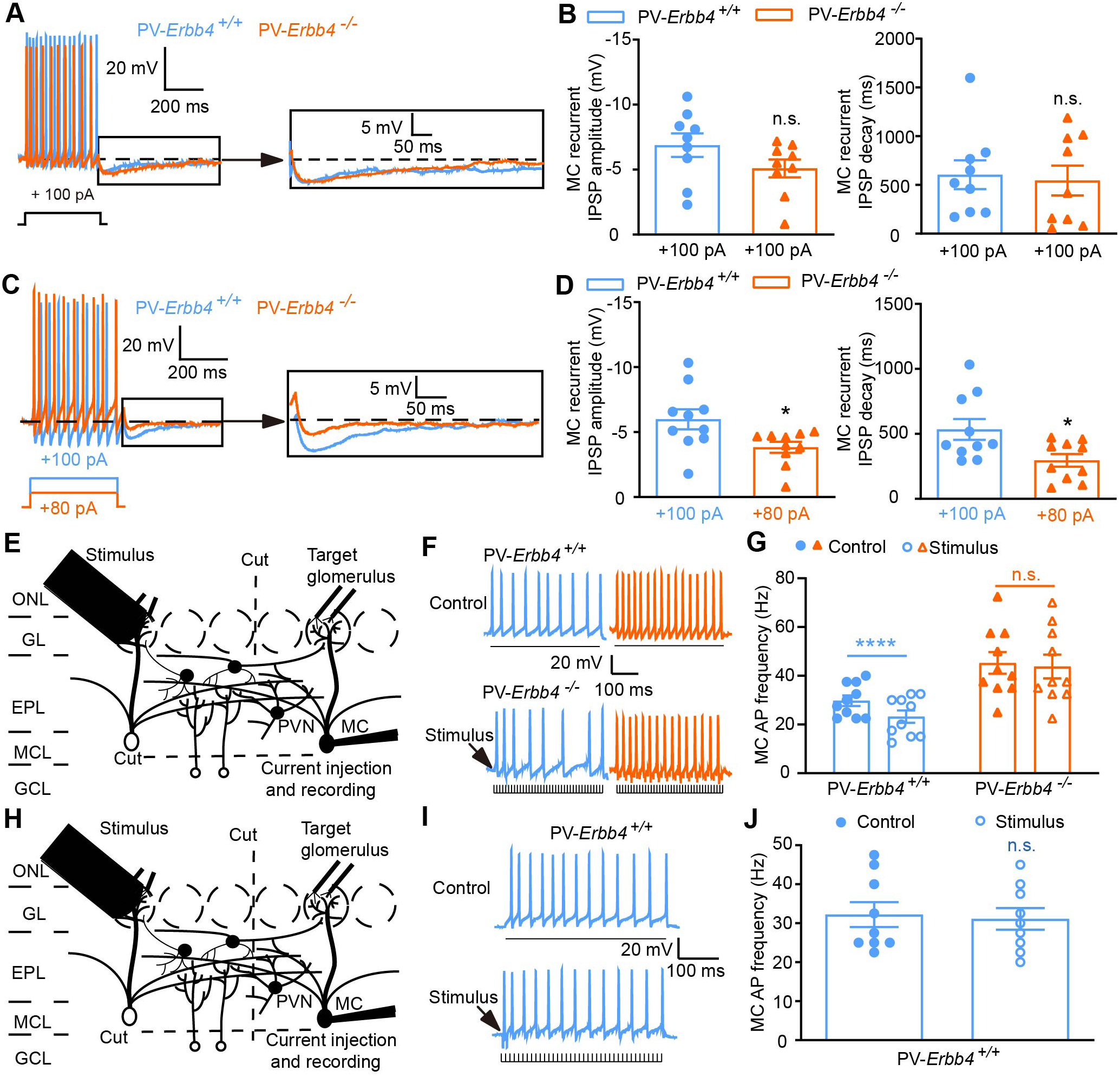
Reduced recurrent inhibition and EPL lateral inhibition of MCs in PV-*Erbb4^−/−^* mice. (A and B) Recurrent inhibition did not increase alongside MC hyperactivity in PV-*Erbb4^−/−^* mice (*n* = 9 from 5 and 6 mice; for peak amplitude, *t_(16)_* = 1.57, *P* = 0.1351; for decay time constant, *t_(16)_* = 0.28, *P* = 0.7821, unpaired *t* test). (C and D) Recurrent IPSPs elicited by the same number of MC APs were smaller in PV-*Erbb4^−/−^* mice. In PV-*Erbb4^−/−^* mice, an 80 pA current induced ten APs, whereas in PV-*Erbb4^+/+^* mice, a 100 pA current was needed to induce ten APs. Both the peak amplitude (*n* = 10 from 6 and 5 mice, *t_(18)_* = 2.46, *P* = 0.0242, unpaired *t* test) and the decay time constant (*t_(18)_* = 2.56, *P* = 0.0198, unpaired *t* test) of IPSPs evoked by the same number of APs were reduced in PV-*Erbb4^−/−^* mice. (E) Schematic of the EPL lateral inhibition experimental configuration. Whole-cell recording of an MC (right side of the schematic) upon stimulus of an adjacent glomerulus (left side of the schematic). A cut was made through the GL and GCL between the sites of conditioning stimulation and target glomerulus to isolate EPL lateral inhibition. (F and G) EPL lateral inhibition was observed in the OB of PV-*Erbb4^+/+^* mice but not PV-*Erbb4^−/−^* mice (*n* = 10 from 7 PV-*Erbb4^+/+^*mice, *F_(1, 18)_* = 26.48, *P* < 0.0001; *n* = 10 from 6 PV-*Erbb4^−/−^* mice, *P* = 0.3426, two-way ANOVA). (H-J) Cutting through the EPL as well abolished the EPL lateral inhibition in PV-*Erbb4^+/+^* mice. (H) Schematic of the experimental configuration. (I and J) Representative and quantitative analysis of spike frequency in PV-*Erbb4^+/+^*mice (*n* = 9 from 5 mice, *t_(8)_* = 0.80, *P* = 0.4468, paired *t* test). EPL, external plexiform layer; GL, glomerular layer; GCL, granule cell layer; MCL, mitral cell layer; ONL, olfactory nerve layer; PVN, PV interneuron. Data are presented as means ± s.e.m. * *P* < 0.05, **** *P* < 0.0001, n.s. = not significant.

In addition, we attempted to isolate broader lateral inhibition between MCs and detect the effects of ErbB4 ablation on it. The recorded MC corresponding to the target glomerulus was visually identified using Alexa 488 dye in a whole-cell patch pipette, as described previously (Hu et al., 2017). A stimulating electrode was then placed into the glomerulus located 3–4 glomeruli (∼400 μm) caudally from the target glomerulus. Before recording, we sectioned the GL and the GCL between the sites of conditioning stimulation and the target glomerulus to eliminate interglomerular and GC-mediated lateral inhibition (Figure 8E). As shown in Figure 8F and 8G, a burst of conditioning stimuli (40 pulses at 100 Hz) caused a reduction in the MC firing rate in PV-*Erbb4^+/+^* mice, but not in PV-*Erbb4^−/−^* mice. When we cut through the EPL, this broader lateral inhibition was completely abolished in PV-*Erbb4^+/+^* mice (Figure 8H-8J), indicating that ErbB4-expressing PV interneurons mediate the broader lateral inhibition.

The above data reveal an important contribution of the PV interneuron ErbB4 to the inhibitory circuit activity in the OB.

### ErbB4 in OB PV interneurons regulates PV-MC inhibitory synaptic transmission and olfactory behaviours

Finally, we evaluated the contribution of PV-ErbB4 in the OB to olfactory behaviour. ErbB4 expression in the PV interneurons in the OB was largely diminished after *loxP*-flanked *Erbb4* mice were stereotactically injected into the OB with AAV-PV-Cre recombinants, but not when injected with control AAV-PV-GFP (Figure 9A-9C).

**Figure 9.**
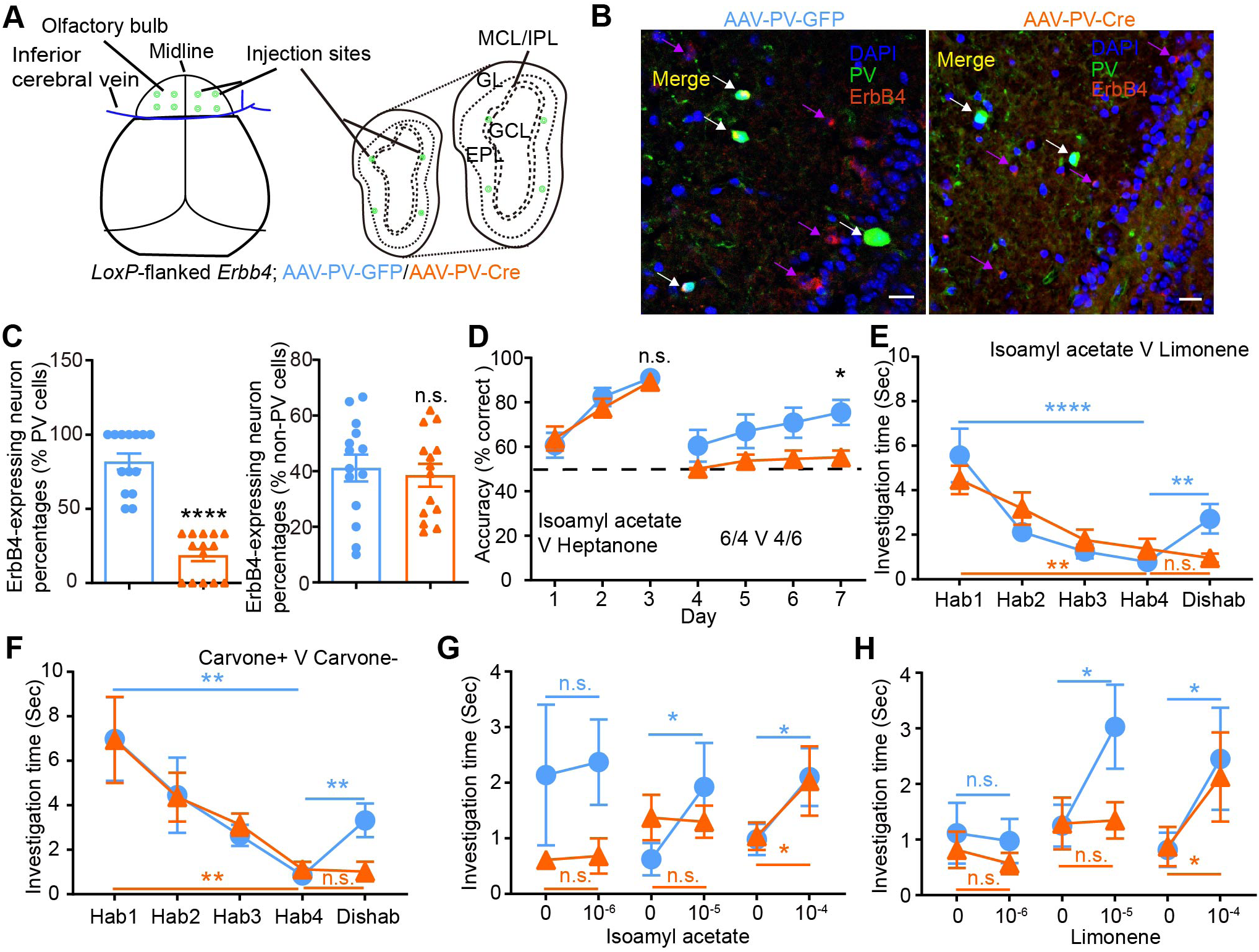
ErbB4 in PV interneurons of the OB is critical for odour discrimination and sensitivity. (A) Schema indicating virus-injection sites. To specifically delete ErbB4 protein in the PV interneurons of the OB, AAV-PV-Cre was injected into the bilateral OB of neonatal *loxP*-flanked *Erbb4* mice. (B and C) Specific deletion of ErbB4 in PV interneurons (white arrow) but not in non-PV cells (purple arrow) of the OB. OB sections from AAV-PV-GFP injected and AAV-PV-Cre injected mice were stained with DAPI, anti-PV and ErbB4 antibody. Scale bars represent 20 μm. *n* = 14 slices from 4 mice per group, *t_(26)_* = 9.30, *P* < 0.0001 for PV interneurons, *t_(26)_* = 0.41, *P* = 0.6833 for non-PV cells, unpaired *t* test. (D) The accuracy in discriminating simple odour pairs was similar for the two groups (*n* = 6 and 7 mice, *F_(1, 11)_* = 0.06, *P* = 0.8147). The accuracy in discriminating difficult odour pairs was significantly lower in AAV-PV-Cre mice (*F_(1, 11)_* = 5.74, *P* = 0.0355, two-way ANOVA). (E) Both animal groups habituated to isoamyl acetate (*n* = 10 mice per group, *F_(3, 54)_* = 20.94, *P* < 0.0001 and = 0.0010, two-way ANOVA). However, the AAV-PV-GFP (*F_(1, 18)_* = 3.95, *P* = 0.0045), but not the AAV-PV-Cre mice (*P* = 0.7107, two-way ANOVA), dishabituated to limonene. (F) Both animal groups habituated to carvone+ (*n* = 10 mice per group, *F_(3, 54)_* = 9.92, *P* = 0.0019 and 0.0035, two-way ANOVA). However, AAV-PV-GFP (*F_(1, 18)_* = 6.72, *P* = 0.0025), but not AAV-PV-Cre mice (*P* = 0.9845, two-way ANOVA), dishabituated to carvone-. (G) AAV-PV-GFP mice were able to detect isoamyl acetate at a concentration of 10^−5^ (*n* = 10 mice, *F_(1, 18)_* = 0.04, 2.63 and 12.42, *P* = 0.8309, 0.0258, and 0.0164 for 10^-6^, 10^-5^, and 10^-4^, two-way ANOVA), whereas AAV-PV-Cre mice only detected isoamyl acetate at a concentration of 10^−4^ (*n* = 10 mice, *P* = 0.9484, 0.8930, and 0.0312, two-way ANOVA). (H) AAV-PV-GFP mice detected limonene at 10^−5^ (*n* = 10 mice, *F_(1, 18)_* = 0.28, 3.81 and 11.79, *P* = 0.7894, 0.0154, and 0.0131 for 10^-6^, 10^-5^, and 10^-4^, two-way ANOVA) but AAV-PV-Cre mice only detected limonene at a higher concentration of 10^−4^ (*n* = 10 mice, *P* = 0.6408, 0.9349, and 0.0496 for 10^-6^, 10^-5^, and 10^-4^, two-way ANOVA). Data are presented as means ± s.e.m. * *P* < 0.05, ** *P* < 0.01, **** *P* < 0.0001, n.s. = not significant. EPL, external plexiform layer; GCL, granule cell layer; GL, glomerular layer; IPL, internal plexiform layer; MCL, mitral cell layer.

ErbB4 reduction in the OB PV interneurons impaired the behavioural discrimination of complex odour mixtures in the reinforced go/no-go task (Figure 9D). Spontaneous dishabituation responses were also impaired in AAV-PV-Cre mice compared with those in the control AAV-PV-GFP group (Figure 9E and 9F). Olfactory detection thresholds for isoamyl acetate and limonene showed a slight increase in AAV-PV-Cre mice compared to AAV-PV-GFP mice (Figure 9G and 9H). Thus, ErbB4 signalling in the PV interneurons in the OB is required for olfactory discrimination and dishabituation in mice.

To directly investigate the effect of ErbB4 signalling on PV-to-MC inhibitory synaptic transmission in OB PV interneurons, we performed optogenetic experiments combined with slice electrophysiology (Figure 9, Supplementary Figure 1A). We stereotaxically injected a 1:1 viral cocktail consisting of AAV-PV-Cre and AAV-EF1α-DIO-hChR2-mCherry into the OB of *loxP*-flanked *Erbb4* mice and wild-type (Wt) control mice to express ChR2 in PV interneurons specifically, and conducted voltage-clamp recordings 10 weeks post-injection. First, we validated reliable ChR2 function in PV interneurons: 473-nm light stimulation (3-s continuous illumination or 10-ms pulses at 1, 5, and 20 Hz) evoked robust photocurrents (Figure 9, Supplementary Figure 1 B), and these responses showed a graded dependence on light intensity (1–3 mW; Figure 9, Supplementary Figure 1C), confirming the effective optogenetic activation of PV interneurons. To verify whether PV interneurons exert inhibitory control over MCs via direct synaptic connections, we performed slice electrophysiological recordings in which PV interneurons were optogenetically activated and postsynaptic currents in MCs were monitored under sequential pharmacological treatments. Application of tetrodotoxin (TTX) completely abolished light-evoked postsynaptic currents in MCs, and subsequent co-administration of TTX and 4-aminopyridine (4-AP, 100 μM) restored these responses, confirming that the recorded currents originated from direct monosynaptic connections between PV interneurons and MCs rather than polysynaptic network activity (Figure 9, Supplementary Figure 1D and 1E). Furthermore, bath application of the GABA_A_ receptor antagonist bicuculline fully blocked light-evoked postsynaptic currents in MCs, demonstrating that synaptic transmission between PV interneurons and MCs is GABAergic (Figure 9, Supplemental Figure 1D and 1E). Collectively, these findings establish that OB PV interneurons form direct inhibitory synaptic connections with MCs, thereby shaping the OB olfactory circuit output. Critically, recordings from postsynaptic MCs revealed that optogenetic activation of PV interneurons induced significantly weaker light-evoked inhibitory postsynaptic currents (oIPSCs) in *loxP*-flanked *Erbb4* mice than in controls, especially across graded light intensities (Figure 9, Supplemental Figure 1F and 1G). Further analysis of the paired-pulse ratio (PPR, 100-ms inter-pulse interval) showed a significant increase in *loxP*-flanked *Erbb4* mice (Figure 9, Supplementary Figure 1H and 1I), suggesting a reduced presynaptic neurotransmitter release probability. Collectively, these optogenetic data demonstrate that ErbB4 in OB PV interneurons supports their inhibitory synaptic output to MCs, which is associated with presynaptic release efficiency.

## Discussion

ErbB4 expression is confined to PV interneurons in the cortex, hippocampus, basal ganglia, and amygdala (Bean et al., 2014; Fazzari et al., 2010) and plays a role in mood (Chen et al., 2022; Chung et al., 2018; Y. Huang et al., 2021) and cognitive behaviours (Cai and Shuman, 2022; Dominguez et al., 2019; Xu et al., 2022). In the present study, we explored the distribution of ErbB4 in OB microcircuits and its functional properties in olfactory behaviour. We found that ErbB4-positive neurones are distributed in all layers of the OB and co-localised with PV interneurons in the IPL and EPL. Furthermore, ErbB4 in PV interneurons regulates OB oscillations and odour-evoked output by maintaining GABA release and lateral and recurrent inhibition of MCs. ErbB4 in the OB as a whole or specifically in the PV interneurons contributes to olfactory behavioural performance. Our findings provide evidence that PV interneurons regulate the activity of MC inhibitory neurotransmission and broad ongoing fluctuations, and hence, the olfactory behaviours of discrimination and dishabituation, via ErbB4 signalling.

We found that ErbB4 in the OB is necessary for odour discrimination and dishabituation. In the EPL and IPL, ErbB4 mRNA and protein are primarily localised to PV interneurons. Our data do not rule out ErbB4 expression in other subtypes of interneurons, as suggested by other studies (Bean et al., 2014; Bovetti et al., 2006; Neddens et al., 2011; Tan et al., 2022). ErbB4 in astrocytes and neural precursors affects the distribution and differentiation of interneurons in mature OB and olfactory abilities (Anton et al., 2004; Moy et al., 2009). ErbB4 regulation of interneuron migration relies on a critical time window (prior to E13.5) (Batista-Brito et al., 2023; Mei and Nave, 2014). The late-onset ErbB4 mutation in the OB may not have been sufficient to cause the developmental phenotypes observed with early gene deletion.

PV interneurons are predominantly excited by Ca^2+^-permeable GluA2-lacking AMPA receptors in the OB and provide strong inhibitory input to MCs (Kato et al., 2013). In the prefrontal cortex, PV interneuron ErbB4 is responsible for the excitatory synaptic plasticity of AMPA and olfactory associative reversal learning (Cai and Shuman, 2022; Xu et al., 2022). Furthermore, transsynaptic binding of ErbB4–Neurexin1β is critical for the activation of ErbB4, extracellular signal-regulated kinase (ERK) 1/2, as well as brain-derived neurotrophic factor (BDNF) in PV interneurons (Cai and Shuman, 2022; Xu et al., 2022). In the OB, Neurexin1β is highly expressed in M/TCs and regulates inhibitory synaptic transmission (Wang et al., 2021). Thus, at the molecular level, the association of ErbB4 with Neurexin1β may facilitate the formation of GluA2-lacking AMPA receptor synapses onto OB PV interneurons, thus regulating GABA release and olfactory processes.

We report here that ErbB4 in PV interneurons most likely regulates the odour detection threshold by reducing background noise and enhancing the SNR of MC responses in the OB. In the neocortex, PV interneurons are implicated in the formation of gamma oscillations, which reduce circuit noise and amplify circuit signals (Sohal et al., 2009). The selective deletion of ErbB4 from fast-spiking PV interneurons increases gamma oscillations in the hippocampus (Del Pino et al., 2013). Our data suggest that ErbB4 in PV interneurons broadly regulates olfactory-related functions by tuning all the frequency bands of ongoing oscillations in the OB. Gamma oscillations in the OB are critical for odour discrimination (Lepousez and Lledo, 2013) and the generation of gamma rhythms in the OB requires inhibitory synaptic transmission in the EPL (Lagier et al., 2004; Lepousez and Lledo, 2013). ErbB4 deficiency in PV interneurons disrupts gamma rhythms by reducing inhibitory synaptic transmission in the OB, leading to the impaired odour discrimination observed in the present study.

The spontaneous activity of MCs is mainly driven by inhibitory interneurons rather than excitatory olfactory sensory neurons (Duchamp-Viret and Duchamp, 1993; Hu et al., 2021; Hu et al., 2017; Stakic et al., 2011). We observed decreases in mIPSC frequency, recurrent inhibition, and lateral inhibition in PV-*Erbb4^−/−^* mice, suggesting that ErbB4 in PV interneurons plays an important role in regulating M/TC activity through recurrent and lateral inhibitory mechanisms. Another interesting finding was that changes in the spontaneous firing rate differed for odour-evoked excitatory and inhibitory responses *in vivo*, suggesting that ErbB4 in PV interneurons dynamically regulates the spontaneous activity of M/TCs in response to different odours. This result reveals a dual regulatory role of ErbB4 in PV interneurons in olfactory coding. For odour-evoked excitatory responses, PV interneurons (via ErbB4) constrain M/TC spontaneous firing at a low baseline, minimising background noise and enhancing the SNR. In contrast, the inhibitory responses maintain a higher spontaneous firing baseline to accentuate odour-evoked suppression. This regulation amplifies the “excitation of excited M/TCs and inhibition of inhibited M/TCs,” a hallmark of the OB’s “winner-take-all” mechanism that sharpens odour identity coding, which is critical for odour discrimination. These data confirm that ErbB4 in PV interneurons does not exert a global modulation, but dynamically tailors M/TC spontaneous activity to odour-evoked responses, thereby optimising the ability of the OB to resolve olfactory cues.

In the present study, we identified a spatially broad (300–400 μm in our experiments) lateral inhibitory circuit in the OB that requires ErbB4 in PV interneurons. Thus, our findings suggest that distinct classes of interneurons mediate interglomerular lateral inhibition in different spatial dimensions. For example, short-axon cells usually have relatively long axonal arbours, spanning as much as 1000 μm (Aungst et al., 2003), that may effectively mediate the broadest (up to 600 μm) spatial dimension of lateral inhibition (Whitesell et al., 2013). In contrast, GCs connect only to their neighbouring MCs (< 50–100 μm apart); their lateral inhibition is narrowly tuned (Miyamichi et al., 2013). Moreover, here we found that PV interneurons mediate spatially moderate (300–400 μm) lateral inhibition via ErbB4. Future studies could benefit from defining the functions (e.g. contrast enhancement) of these distinct spatial dimensions of lateral inhibition in odour information processing. Both recurrent and lateral inhibition implement a “winner-takes-all” mechanism: once the strongest input excites the principal cells, firing in the remaining inputs is inhibited (de Almeida et al., 2009; Espinoza et al., 2018; Karmakar and Sarkar, 2013). The regulation of recurrent and lateral inhibition by ErbB4 sheds light on how PV interneurons mediate this “winner-takes-all” inhibitory microcircuit.

In recent decades, impaired odour discrimination has been observed in the earliest stages of neurodegenerative disorders, such as Alzheimer’s and Parkinson’s diseases (Dan et al., 2021). Olfactory discrimination deficits develop prior to neuropsychological characteristics in patients with Alzheimer’s disease and have significant predictive value for future cognitive decline in healthy aging individuals (Carnemolla et al., 2020; Sohrabi et al., 2012; Uchida et al., 2020; Yoshitake et al., 2022). *ErbB4* has been independently identified as a susceptibility gene for Alzheimer’s disease and schizophrenia (Mei and Nave, 2014; Swaminathan et al., 2012). Here, we showed that the selective loss of ErbB4 in OB PV interneurons impairs olfactory behaviour. Although ongoing fluctuations predict behavioural alterations in health and disease, the contributions of their different frequency bands remain poorly understood (Iemi et al., 2022). As referenced by Iemi et al. (2022), “fluctuation status” herein refers to the spontaneous, endogenous fluctuations in neural activity that occur in the brain prior to sensory stimulation (i.e. pre-stimulus) or in the absence of external sensory input. External cues do not drive this state but arise from intrinsic neural dynamics, and it is primarily characterised by changes in the power of neural oscillations across multiple frequency bands (e.g., alpha, beta, and their neighbouring frequencies, collectively termed “alpha+ band” in Iemi et al. (2022)). Critically, this fluctuation status is distinct from stimulus-evoked neural responses; it reflects a pre-existing “baseline” state of the neural system that modulates subsequent sensory processing and behaviour. Consistent with Iemi et al. (2022), previous work has established that such ongoing neural fluctuations strongly predict behavioural variability in both health (e.g. slower reaction times and reduced perceptual accuracy) and disease (e.g. age-related cognitive decline and schizophrenia) by regulating neuronal excitability. In our study, we extended this concept to olfactory function, focusing on “broad ongoing fluctuations” (encompassing multiple frequency bands) and linking their dynamics to OB output. Specifically, these fluctuations influence the ability of the OB to support rapid recovery from olfactory adaptation and to mediate complex odour discrimination, thereby uncovering a novel role for endogenous neural fluctuation status in olfactory circuit function. This study links the broad, ongoing fluctuations to OB output and, hence, controls rapid recovery from olfactory adaptation and complex odour discrimination. Our findings will help us better understand the molecular and cellular mechanisms underlying early olfactory dysfunction in neurodegenerative diseases and schizophrenia and provide a potential strategy for treatment.

In summary, our study demonstrates the crucial role of ErbB4 in olfactory behaviours involving ErbB4-expressing PV interneurons. Our results are most likely attributable to presynaptic impairments in inhibitory microcircuits of the OB, which affect odour processing information and lead to abnormal output signals from the M/TCs. The data presented may be promising not only for understanding the physiological functions of ErbB4 in the OB, but also for providing early diagnosis of Alzheimer’s disease and schizophrenia and potential therapeutic applications, particularly in patients with olfactory disorders.

## Materials and methods

### Reagents and animals

Rabbit polyclonal anti-ErbB4 antibody (sc-283; Santa Cruz Biotechnology, Dallas, TX, USA) was used at dilutions of 1:2,000 for immunoblotting and 1:100 for immunostaining. Rabbit polyclonal anti-ErbB4 antibody (19943-1-AP; Proteintech, Rosemont, IL, USA) was diluted 1:100 for immunostaining, while rabbit monoclonal anti-ErbB4 antibody (#4795; Cell Signalling Technology, Danvers, MA, USA) was used at 1:1000 for immunoblotting. Mouse monoclonal anti-phosphotyrosine antibody (05-321; Sigma-Aldrich, St. Louis, MO, USA) was applied at 0.5 μg per 200 μg total protein for immunoprecipitation, and mouse monoclonal anti-parvalbumin (PV) antibody (P3088; Sigma-Aldrich) was diluted 1:7000 for immunostaining. Anti-β-actin antibody (#4970; Cell Signalling Technology) served as the loading control at 1:5000 dilution for immunoblotting. Goat anti-rabbit IgG conjugated to Alexa Fluor 488 (A11089; Invitrogen, Waltham, MA, USA) and goat anti-mouse IgG conjugated to Alexa Fluor 594 (A11037; Invitrogen) secondary antibodies were both used at 1:400 for immunostaining. The sodium channel blocker tetrodotoxin (TTX, 1069; Tocris, Bristol, United Kingdom) and GABA_A_ receptor antagonist (+)-bicuculline (ALX-550-515; Enzo Life Sciences, Farmingdale, NY, USA) were obtained from the listed suppliers. All remaining chemicals were purchased from Merck (Sigma– Aldrich).

PV-*Erbb4*^−/−^ were generated by crossing PV-Cre mice with *loxP*-flanked *Erbb4* mice, while PV-*Erbb4^+/+^* mice carried the PV-Cre transgene and the Wt *Erbb4* allele (Garcia-Rivello et al., 2005; Wen et al., 2010; Xu et al., 2022). The mice were housed under a 12-h light/dark cycle and had ad libitum access to water and food, except during the behavioural tests. All experiments were performed on both sexes of mice. All experimental procedures were conducted in accordance with the guidelines described in the revised Regulations for the Administration of Affairs Concerning Experimental Animals (2011) in China, and were approved by the local Institutional Animal Care and Use Committee.

### Immunoprecipitation

Protein samples were resuspended in immunoprecipitation (IP) buffer supplemented with 50 mmol/L HEPES, 150 mmol/L NaCl, 1 mmol/L ZnCl₂, 1.5 mmol/L MgCl₂, 10% glycerol, 1% Triton X-100, 0.5% Nonidet P-40, and phosphatase/protease inhibitors (Xu et al., 2022). The samples were then incubated with the appropriate primary antibodies at 4 °C overnight, followed by incubation with Protein A Sepharose CL-4B beads (GE Healthcare Biosciences, Uppsala, Sweden) for 2 h at 4 °C with gentle rotation. Immune complexes were collected by centrifugation, and after three sequential washes with IP buffer, bead-bound proteins were eluted and subjected to immunoblotting.

### Western blot analysis

Tissues were homogenised in the homogenisation buffer containing 50 mM Mops (pH 7.4), 320 mM sucrose, 100 mM KCl, 0.5 mM MgCl_2_, 0.2 mM dithiothreitol, and phosphatase and protease inhibitors. Homogenates were resolved by 7.5% SDS/PAGE and transferred to nitrocellulose membranes (Millipore, Billerica, MA, USA), which were blocked with 3% bovine serum albumin (BSA) in Washing buffer (100 mM NaCl, 0.1% Tween-100, 10 mM Tris, pH 7.5). Membranes were incubated overnight with the primary antibody in 1% BSA and then probed with the secondary antibody for 1 h. Signals were visualised using the Immobilon Western Chemiluminescent horseradish peroxidas substrate. Immunoblotting was performed using ImageJ software (National Institutes of Health, Bethesda, MD, USA).

### Virus injections

Viral injections were performed as described previously (Kato et al., 2013). Neonatal *loxP*-flanked *Erbb4* mice (0–3 days old) were cryoanesthetized in an ice-cold chamber and head-fixed using a stereotaxic device (RWD68513; RWD Life Science, Shenzhen, China). Viruses driven by the CAG promoter, including pAOV-CAG-EGFP-T2A-Cre (catalog No. CN396, serotype AAV2/8) and pAOV-CAG-EGFP-2A (catalog No. CN394, serotype AAV2/8) were purchased from Obio Technology, Shanghai, China. Additional recombinant adeno-associated viruses, namely rAAV-fPV-CRE-bGH polyA (PT-0275), rAAV-PV-EGFP-bGH polyA (PT-0154) and rAAV-EF1α-DIO-hChR2(H134R)-mCherry-WPRE-hGH polyA (PT-0002, all serotype AAV2/9), were obtained from BrainVTA, Wuhan, China. And the titre was 10^12^ genome copies/mL. For optogenetic activation of OB PV interneurons, a viral cocktail (1:1) of AAV-PV-CRE and AAV-EF1α-DIO-hChR2-mCherry was stereotactically injected into the OB of *loxP*-flanked *Erbb4* or control mice. The injection coordinates were as follows (anteroposterior, mediolateral, and dorsoventral relative to the intersection of the midline and the inferior cerebral vein, in mm): (+0.2, ±0.3, − 0.9/0.7), (+0.2, ±0.6, −0.9/0.7), (+0.5, ±0.3, −0.7/0.5) and (+0.5, ±0.6, −0.7/0.5). The virus (20 nL per site) was injected at a rate of 23 nL/s using a glass pipette. Behavioural tests were performed 10 weeks after the virus injection.

### Behavioural analysis

Behavioural analysis was performed by investigators blinded to the experimental animals’ genotypes.

### Go/no-go test

Mice were deprived of water, allowed to explore the setup (Thinkerbiotech, Nanjing, China), and learned to lick a metal tube just below the odour port to obtain water, as previously described (Hu et al., 2021). The mice self-initiated the trials by poking their noses into the sampling port, which interrupted the infrared light beam. All odours were diluted to 0.01% in mineral oil and further diluted to 1:20 in air using olfactometers. Odour pairs were presented for 2 s in randomised order 0.5 s after trial initiation. Prior to training on the go/no-go task, the mice were trained on the go/go task for 2 days. During the go/go task, if the mice licked within the odour presentation period when presented with either of the odour pairs, they received a water reward. The go/no-go task was then performed to train the mice to discriminate between a pair of odours in order to receive the water reward. Mice learned to lick the metal tube to receive water in response to the rewarded odour (hit) at the end of the trial (2.5-s total trial duration) and to avoid licking the metal tube for the unrewarded odour (correct rejection). Licking when presented with the unrewarded odour (false alarm) led to no water reward and a timeout of up to 10 s. Performance “accuracy” was calculated as the percentage of correct trials (number of hits and correct rejections) over the total number of trials. The weight of the mice under water restriction was strictly monitored (>85% of the initial body weight) during the entire test. The eight-channel semiautomated olfactometers, data acquisition, and analysis were controlled using computer programs written in LabVIEW and MATLAB.

### Habituation/dishabituation test

Odours were presented by placing a cotton swab scented with an odour solution (150 μL, 1:2000, freshly prepared with mineral oil before each experiment) above the floor of the animal’s home cage, as previously described (Hu et al., 2021). The mice were familiarised with the first odour in four successive trials (habituation) and then exposed to a novel odour in the fifth trial (dishabituation). Five successive 2-min trials were separated by 2 min. Each trial was videotaped and the swab investigation time was quantified offline. Sniffing behaviour was defined as the time when the mouse had its nose within a 1-cm radius from the surface of the cotton swab. A lack of novel odour discrimination was considered to occur when mice spent as much time investigating the swab during the dishabituation trial as during the fourth habituation trial.

### Odour detection threshold test

Odour detection thresholds were obtained according to previously described procedures (Nicolis di Robilant et al., 2019). Odours (isoamyl acetate and limonene) were freshly diluted with mineral oil to different concentrations (10^-6^, 10^-5^, and 10^-4^ in 150 μL) and applied to a cotton swab, then placed above the floor of the animal’s cage along with mineral oil on the other side. Each trial lasted 2 min with 2-min intertrial intervals. Mice were deemed able to detect the odour when they spent more time sniffing the diluted odour than the mineral oil, and the minimum concentration detected by the mice was taken as the threshold.

### Buried food pellet test

All chow pellets were removed from the home cages 24 h before testing. The mice had ad libitum access to water. Before each experimental trial, 10-to 12-week-old mice were familiarised with the test cage for 10 min. In each trial, a single mouse was placed in the test cage (46 × 30 × 16 cm) to recover a 0.2-g food pellet buried 0.5 cm below the bedding material surface. Food pellet locations were randomly selected. The time between when the mouse was placed in the cage and when it grasped the food pellet with its forepaws or teeth was recorded and defined as latency. The mice were allowed to consume food pellets and were returned to their cages. The food pellet was presented to the mouse for consumption if it was not found within 300 s, in which case, the latency was recorded as 300 s. A visible food pellet test was conducted as a control. The procedures were identical, except that the pellets were placed randomly on the bedding surface.

### RNAscope

To examine the expression of the *Erbb4* gene, RNAscope was performed using an RNAscope Fluorescent Multiplex Kit (Advanced Cell Diagnostics (ACDBio), Newark, CA, USA) according to the manufacturer’s recommendations. Briefly, the OB was cut into 20-µm coronal sections. Sections were serially thaw-mounted onto 20 slides (Fisherbrand, Waltham, MA, USA) through the entire OB and then air-dried for 1 h at room temperature prior to storage at −80 °C. RNAscope probes were obtained from ACDBio for *Erbb4* (Cat #318721) and *Pv* (Cat #421931-C2). The sections were counterstained with the nuclear marker DAPI (Cat #104139, Abcam, Cambridge, United Kingdom). *Erbb4* mRNA intensity in the OB was measured using ZEN Blue 3.0 and ImageJ software. Experimenters blinded to the group assignments performed image collection and subsequent data analysis.

### Immunohistochemistry

Mice (4 weeks old) were perfused transcardially with 4% paraformaldehyde (PFA). Dissected brains were postfixed overnight and dehydrated in 30% sucrose for 24 h at 4 °C, after which the OBs were cut into 40-μm-thick horizontal sections on a vibratome (Leica Microsystems, Wetzlar, Germany). OB sections were processed for double immunofluorescence in 0.1 M PBS (pH 7.4) as follows: three washes in 0.1 M PBS, 10 min each; blocking in 0.1 M PBS containing 10% normal goat serum, 1% BSA, and 0.25% Triton X-100 overnight at 4 °C; primary antibody incubation for 40 h at 4 °C in blocking solution; three washes; secondary antibody for 90 min at room temperature in blocking solution; three washes; rinsing with PBS and mounting with a mounting medium (Vector Laboratories, Burlingame, CA, USA). Immunoreactivity was assessed using Alexa Fluor 488– and 594-conjugated goat anti-mouse IgG antibodies. Sections were analysed using a confocal microscope (Zeiss LSM710, Carl Zeiss AG, Oberkochen, Germany) at 20× and 60× magnifications. ErbB4 intensity in the OB was measured using ZEN Blue 3.0 and ImageJ software. Cresyl violet staining was performed as described previously. The optimal cutting temperature compound-embedded OBs were cut into 20-µm sections. OB sections were stained with 0.1% cresyl violet for 35 min, dehydrated using ascending grades of alcohol, cleared with xylene, mounted, and observed under an Olympus microscope.

### Slice preparation and patch-clamp electrophysiology

Acute OB slices were prepared as described previously (Hu et al., 2021; Hu et al., 2017). Briefly, animals were heavily anaesthetised with ketamine/xylazine (140/20 mg/kg, intraperitoneally) and decapitated. The OB was quickly removed and immersed in ice-cold, and oxygenated (95% O_2_/5% CO_2_) ACSF containing 124 mM NaCl, 3 mM KCl, 2 mM CaCl_2_ 1.3 mM MgCl_2_, 25 mM NaHCO_3_, 1.25 mM NaH_2_PO_4_, and 10 mM glucose. Horizontal slices (300 μm) were cut with a VT1000s vibratome (Leica), recovered at 37 °C for 30 min and then maintained at 25 °C.

OB slices were transferred to a recording chamber, which was warmed to 33 °C and superfused with oxygenated ACSF (2 mL/min). Neurones were visualised with infrared optics using an upright microscope equipped with a 60× water-immersion lens (BX51WI; Olympus, Tokyo, Japan) and an infrared-sensitive charge-coupled device camera. MCs and PV interneurons were identified based on their fluorescence, morphology, size, location, and electrophysiological characteristics.

For cell-attached mode, recording pipettes with resistances of 6–8 MΩ were pulled from borosilicate glass capillaries (P97; Sutter Instrument Co., Novato, CA, USA) and filled with ACSF. To record olfactory-nerve-evoked responses, monophasic square pulses (200 μs) with an intensity of 0.2 mA or gradual increasing intensities of 0.1, 0.15, 0.2, 0.25, and 0.3 mA were delivered through a concentric electrode that was placed on the outermost layer of the OB, near the recorded MC. The firing frequency was defined as the number of action potentials/time windows. A 5-s window prior to olfactory nerve stimulation was used to assess the spontaneous firing rate, and the first 1-s window after stimulation of the olfactory nerve was used to assess the evoked response. The signal-to-noise ratio (SNR) was defined as the ratio of evoked responses to spontaneous firing 5 s before the olfactory nerve stimulation.

To record whole-cell action potentials and mEPSCs, pipettes were filled with a solution containing 140 mM K-methylsulfate, 4 mM NaCl, 10 mM HEPES, 0.2 mM EGTA, 4 mM MgATP, 0.3 mM Na_3_GTP, and 10 mM phosphocreatine (pH was adjusted to 7.4 with KOH). MCs were held at −65 mV in the presence of 20 µM bicuculline and 1 μM TTX during recording of mEPSCs.

To isolate mIPSCs, MCs were held at −65 mV in the presence of 50 µM AP5, 20 µM NBQX and 1 μM TTX with a CsCl-base intracellular solution containing (mM): 135 CsCl, 10 HEPES, 0.2 EGTA, 2 Na_2_ATP, 0.3 Na_3_GTP, and 10 glucose.

Lateral inhibition recordings were performed as previously described (Arevian et al., 2008; Hu et al., 2017; Whitesell et al., 2013). Briefly, a concentric bipolar stimulating electrode was placed into the caudal glomerulus located 3–4 glomeruli (∼400 μm) caudally from the one correlated with the recorded MC. The intensity of the stimulation was moderate (100 μA) to avoid stimulating the glomerulus of the recorded MC, which would invariably evoke excitatory postsynaptic potentials (EPSPs). Pulses of 200-μs duration were delivered at 100 Hz and synchronised with the recording using a Master-8 stimulator (A.M.P.I., Jerusalem, Israel). Alexa 488 dye (100 μM) was added to the intracellular solution to visualise cell morphology.

Photostimulation (273 nm, 10-ms pulses) of ChR2-expressing PV interneurons was delivered through an optical fibre placed near the slice and connected to an intelligent optogenetic system (Aurora-220; NEWDOON, Hangzhou, China). Photostimulation was triggered using Clampex software (Molecular Devices, San Jose, CA, USA). During optogenetic experiments, PV interneurons were voltage-clamped at –70 mV, while MCs were maintained at a holding potential of 0 mV.

Signals were acquired with a MultiClamp 700 B amplifier (Molecular Devices), filtered at 2 kHz, and sampled at 10 kHz using a Digidata 1440A interface (Molecular Devices) and Clampex 10.2 software (Molecular Devices). The data were accepted when the series resistance remained within 15% of the initial value. Miniature events in a 5-min recording period were analysed using the Mini Analysis software (version 6.07; Synaptosoft, Decatur, GA, USA).

### Single-cell RT-PCR

At the end of the recording session, gentle suction was applied to aspirate the cytoplasm into the pipette, while maintaining a tight seal. After complete incorporation of the soma, the pipette was carefully removed from the neuron to perform an outside-out patch recording, and then quickly removed from the bath. The harvested contents were transferred into PCR tubes. RT was performed in a final volume of 20 μL using a QuantiTect Reverse Transcription Kit (Cat. No. 205311; QIAGEN, Venlo, The Netherlands). Next, outer and nested primer sets were used for two consecutive rounds of PCR as described previously (Vullhorst et al., 2009). For the first round of PCR, all targets were amplified simultaneously using outer primer pairs for PV and ErbB4 (first set PCR primer sequences: GCCTGAAGAAAAAGAACCCG and AATCTTGCCGTCCCCATCCT for PV, CCAGCCCAGCGCTTCTCAGTCAG and GTATTTCGGTCAGGTTCTTTAATCC for ErbB4, 20 pmol of each) in a final volume of 100 μL for 21 cycles. Each cycle comprised a 94 °C denaturation for 30 s, 60 °C annealing for 30 s, and 72 °C extension for 30 s. For the second round of PCR, PV and ErbB4 were separately reamplified using specific nested primer sets for 35 cycles of PCR, as described above, with 2 μL of the first PCR product as template (second primer sequences: CGGATGAGGTGAAGAAGGTGT and TCCCCATCCTTGTCTCCAGC for PV, CTGACCTGGAACAGCAGTACCGA and AGGCATAGCGATCTTCATATAGT for ErbB4). The products were assayed on 2% agarose gels stained with Gel Red, with dl1000 as the molecular weight marker.

### Implantation of electrodes for in vivo electrophysiological recordings

After deep anaesthesia, the mice were secured in a stereotaxic frame. The fur on the scalp surface was removed from the midpoint between the ears and the midline of the orbits. For the spike and LFP recordings, tetrodes (single-wire diameter, item no. PF000591, RO-800, 0.0005”/12.7 μm, coating 1/4 hard PAC, Sandvik AB, Stockholm, Sweden) were implanted into the hole drilled above the OB (4.28 mm anterior from bregma, 1.0 mm lateral from the midline). The electrodes were inserted into the brain after the dura mater was punctured. Each electrode was connected to a 16-channel electrode interface board (EIB-16; Neuralynx, Bozeman, MT, USA). Signals were sent to a headstage, amplified using a 16-channel amplifier (Plexon Inc., Dallas, TX, USA), and monitored in real time to ensure optimal placement within the ventral MC layer. Despite the presence of diverse cell types in the OB, only the spiking activity of M/TCs can be captured via tetrode recordings (Kay and Laurent, 1999; Doucette et al., 2011). A stainless steel screw was inserted into the parietal bone (1 mm posterior to the bregma and 1 mm from the midline) and connected to the ground as the reference electrode. Finally, a custom-designed aluminium head plate was sealed to the skull surface using dental cement to enable head fixation during recording. The mice were returned to their home cages for at least a week.

### Electrophysiological recordings in awake mice

Recordings began at least one week after electrode implantation. The mice were head-fixed with two horizontal bars fixed to the headplate using two screws, and were able to walk on an air-supported floating polystyrene foam ball (Thinkerbiotech). For spike recordings, the signals from the electrodes were filtered at 300–5000 Hz and sampled at 40 kHz. Odour stimulus event markers were recorded alongside the spike recording. For LFP recordings, the signals were amplified by a 2000× gain, bandpass filtered (0.1–300 Hz, and sampled at 1 kHz (Plexon Inc.).

### Odour application for electrophysiological recordings in awake mice

A standardised panel of eight odours (isoamyl acetate, 2-heptanone, phenyl acetate, benzaldehyde, dimethylbutyric acid, n-heptane acid, n-pentanol, and 2-pentanone; purchased from Sinopharm Chemical Reagent Co., Ltd., Beijing, China) was prepared using an odour delivery system (Thinkerbiotech). Charcoal-filtered air was supplied to the mouse nose at a constant rate of 1 L/min to eliminate airflow interference. This method has been validated in previous studies to effectively limit odour stimulation to within 2 s. All odours were dissolved in a 1% (v/v) dilution of mineral oil. During odour stimulation, a stream of charcoal-filtered air flowed over the oil at a total flow rate of 1 L/min, and the vapour-phase concentration was further diluted by the carrier airflow to 1/20. Mice were first anaesthetised with isoflurane for approximately 20 s and then rapidly transferred to a polystyrene foam ball. Before the start of *in vivo* electrophysiological recordings, the mice were given 10 min to acclimate to the fixation and recover from isoflurane. Odour presentation was controlled using an odour delivery system coupled to a solenoid valve driven by a digital-to-analogue converter. Each odour was presented for 2 s, with an inter-trial interval of approximately 24 s.

### Offline spike sorting and statistics of the unit data

Spikes were identified and sorted from the raw data using Offline Sorter V4 software (Plexon Inc.). Spikes were detected and sorted when they had an amplitude exceeded 5 times the noise standard deviation. Principal component analysis of the data was performed to separate the different units. When <0.75% of the inter-spike intervals were <1 ms, the unit was classified as a single unit. Data from 4 s before to 6 s after the onset of the odour stimulation event were extracted, and the spike firing rate was averaged within 100-ms bins to generate a peristimulus time histogram (PSTH). The spontaneous and odour-evoked firing rates were calculated by averaging the spikes recorded during 4–2 s before odour stimulation and 2 s after the start of odour stimulation, respectively. To determine whether the odour evoked a significant response, we used the Wilcoxon signed-rank test to compare the spontaneous (baseline) firing rate with the odour-evoked firing rate across all trials. Unit-odour pairs (*P* > 0.05) were defined as non-responsive; those (*P* < 0.05) were defined as responsive, and responsive pairs were further categorised as excitatory (odour-evoked firing rate > spontaneous firing rate) or inhibitory (odour-evoked firing rate < spontaneous firing rate). Subtracting the baseline firing rate from the odour-evoked firing rate yielded the mean firing rate (MFR). The SNR was defined as the odour-evoked firing rate divided by the baseline firing rate.

### LFP signal analysis

The LFP signals were analysed using a MATLAB script and divided into different frequency bands: theta (2–12 Hz), beta (15–35 Hz), low-gamma (36–65 Hz), and high-gamma (66–95 Hz). The spectral power of each frequency resolution was calculated over a 4-s window. The spectral powers of all frequencies within the bandwidth were averaged.

### Statistical analysis

Data are presented as the mean ± standard error of the mean (SEM). Unpaired or paired *t* tests were used to compare data between the two groups. The Chi-square test was used to compare the two constituent ratios. One-way or two-way ANOVAs were used to comparemore than two groups (western blot, behavioural, and electrophysiological studies). Tukey’s post-hoc adjustment was used for multiple pairwise comparisons in the behavioural tests. All tests were two-sided, and statistical significance was set at *P* < 0.05. Statistical analyses were performed using the GraphPad Prism software.

## Additional information

### Competing interest

The authors declare no competing interests.

## Funding

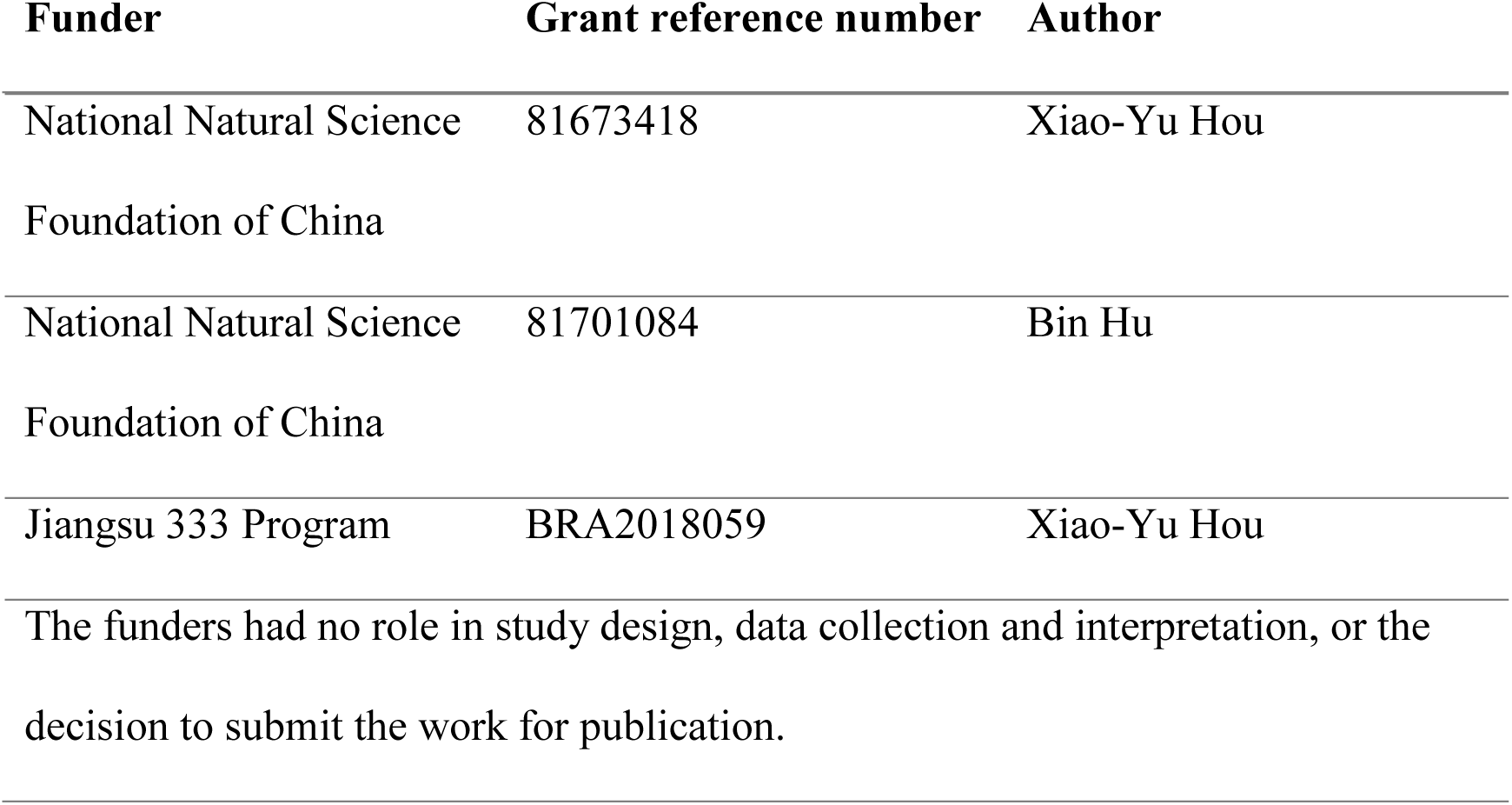

## Author Contributions

Bin Hu, Data curation, Formal analysis, Funding acquisition, Investigation, Methodology, Writing—original draft; Chi Geng, Data curation, Formal analysis, Investigation, Methodology, Writing—original draft; Feng Guo, Ying Liu, Data curation, Formal analysis, Investigation, Methodology; Ran Wang, Investigation, Methodology; You-Ting Chen, Methodology; Xiao-Yu Hou, Conceptualization, Resources, Supervision, Funding acquisition, Validation, Project administration, Writing—review and editing.

## Ethics

All experimental procedures were conducted in accordance with the guidelines described in the revised Regulations for the Administration of Affairs Concerning Experimental Animals (2011) in China and approved by the local Institutional Animal Care and Use Committee.

## Data availability

All data generated or analyzed during this study are included in the manuscript and supporting files.

## Figure legends

**Figure 1 and 2—figure supplement 1.** Uncropped western blot images. (A) Uncropped western blot images for p-ErbB4 in Figure 1C. (B) Uncropped western blot images for total ErbB4 in Figure 1C. (C) Uncropped western blot images for ErbB4 in Figure 1F. (D) Uncropped western blot images for ErbB4 in Figure 2K.

**Figure 2—figure supplement 2.** The overall lamina structures of the OB were not affected in PV-*Erbb4^−/−^* mice. Cresyl violet staining was performed to examine coronal structure sections of the OB at 2 and 6 months of age. Scale bars represent 200 μm for the overall views and 50 μm for the magnified images. EPL, external plexiform layer; GCL, granule cell layer; GL, glomerular layer; IPL, internal plexiform layer; MCL, mitral cell layer; ONL, olfactory nerve layer.

**Figure 4—figure supplement 1.** Analysis of M/TC spike waveform features in PV-*Erbb4^+/+^* and PV-*Erbb4^−/−^* mice. (A) Separation of the 2 units (red and blue dots) using PCA. (B) Examples of isolated two different single units (unit a, red; unit b, blue). (C) Examples of spike waveform shapes of M/TCs in PV-*Erbb4^+/+^* and PV-*Erbb4^−/−^* mice. (D) Quantitative analysis of peak-to-valley ratio of spikes (*t_(52)_* = 1.92, *P* = 0.0610, unpaired *t* test). (E) Quantitative analysis of half-valley width of spikes (*t_(52)_* = 1.41, *P* = 0.1648, unpaired *t* test). Data are presented as means ± s.e.m. n.s. = not significant.

**Figure 7—figure supplement 1.** Electrophysiological properties of OB PV interneurons in PV-*Erbb4^+/+^* and PV-*Erbb4^−/−^* mice. (A and B) Reduced frequency and amplitude of OB PV interneuron mEPSCs in PV-*Erbb4^−/−^*mice (*n* = 6 from 3 PV-*Erbb4^+/+^*mice, *n* = 4 from 3 PV-*Erbb4^−/−^* mice; for frequency, *t_(8)_* = 2.43, *P* = 0.0411; for amplitude, *t_(8)_* = 3.15, *P* = 0.0136, unpaired *t* test). (C) Increased frequency of OB PV interneuron APs in PV-*Erbb4^−/−^* mice (*t_(8)_* = 2.394, *P* = 0.0436, unpaired *t* test). (D) Quantitative analysis of AP amplitude in OB PV interneurons (*t_(8)_* = 1.52, *P* = 0.1660, unpaired *t* test). (E) Quantitative analysis of AP half-width in OB PV interneurons (*t_(8)_* = 0.99, *P* = 0.3528, unpaired *t* test). Data are presented as means ± s.e.m. * *P* < 0.05, **** *P* < 0.0001, n.s. = not significant.

**Figure 8—figure supplement 1.** Approximately the same AP firing rate elicited by different current intensities across PV-*Erbb4^+/+^* and PV-*Erbb4^−/−^* mice. (A) A current injection of ∼ 80 pA in PV-*Erbb4^−/−^* mice elicited approximately the same firing rate as a current injection of ∼ 100 pA in PV-*Erbb4^+/+^* mice (*n* = 10 from 6 and 5 mice, *t_(18)_* = 0.09, *P* = 0.9316, unpaired *t* test). Data are presented as means ± s.e.m. n.s. = not significant.

**Figure 9—figure supplement 1.** ErbB4 deletion in OB PV interneurons attenuates PV interneuron-mediated inhibitory modulation of MCs. (A) Schematic diagram illustrating the experimental paradigm, including stereotaxic virus injection and subsequent slice electrophysiological recordings; blue light illumination was delivered to OB slices, with recordings performed from PV interneurons expressing ChR2 or postsynaptic MCs. (B) Representative voltage-clamp recordings of PV interneuron currents in response to 3-s continuous 473-nm light illumination and pulsed light stimulation (10-ms pulses at 1, 5, and 20 Hz). (C) Representative voltage-clamp traces of PV interneuron currents evoked by graded 473-nm light intensities (1, 2, and 3 mW). (D and E) Representative traces and quantitative analysis of oIPSCs in MCs under distinct pharmacological conditions (Con, TTX, TTX + 4-AP, and bicuculline (Bic)). Statistical comparisons: Con vs. TTX, *P* = 0.0004; TTX vs. TTX + 4-AP, *P* < 0.0001; TTX + 4-AP vs. Bic, *P* < 0.0001 (n = 3 from 3 Wt mice; F*_(3, 8)_* = 47.78, one-way ANOVA). (F and G) Representative traces and quantitative analysis of light-evoked MC oIPSCs across graded illumination intensities in Wt and *loxP*-flanked *Erbb4* mice (n = 10 from 3 mice per group; two-way ANOVA, F*_(1, 53)_* = 23.26, *P* < 0.0001). (H and I) Representative traces and quantitative analysis of PPR (100-ms inter-pulse interval) of light-evoked MC oIPSCs in Wt and *loxP*-flanked *Erbb4* mice (n = 10 from 3 mice per group; *P* = 0.0163, t*_(18)_* = 2.651, unpaired t test). Data are presented as means ± s.e.m. * *P* < 0.05, *** *P* < 0.001, **** *P* < 0.0001. EPL, external plexiform layer; GCL, granule cell layer; GL, glomerular layer; IPL, internal plexiform layer; MCL, mitral cell layer; ONL, olfactory nerve layer; PVN, parvalbumin interneuron.

## Notes

### Competing Interest Statement

The authors have declared no competing interest.

### Summary of Updates

All figures revised; Data quantified; Discussion revised to discuss the significance of two key findings; Materials and methods section refined; optogenetic experiments added; Supplemental files updated.

### Funding

- National Natural Science Foundation of China: 81673418, 81701084
- Jiangsu 333 Program: BRA2018059

