## Supplementary figures and images for "Parvalbumin interneuron ErbB4 controls ongoing network oscillations and olfactory behaviours in mice"

### Figure1 and 2 and supplement figure 1

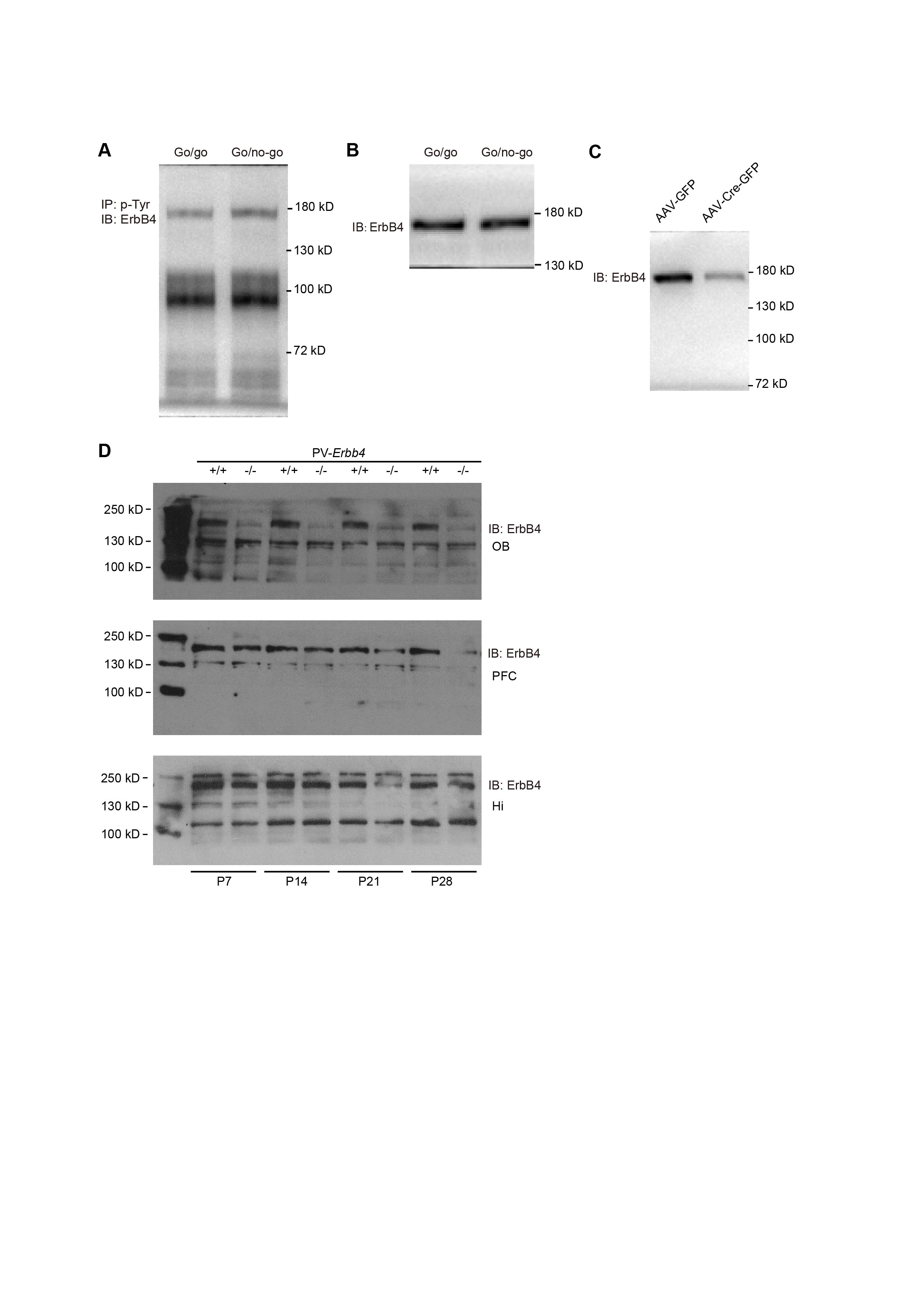

### Figure2 and supplement figure 2

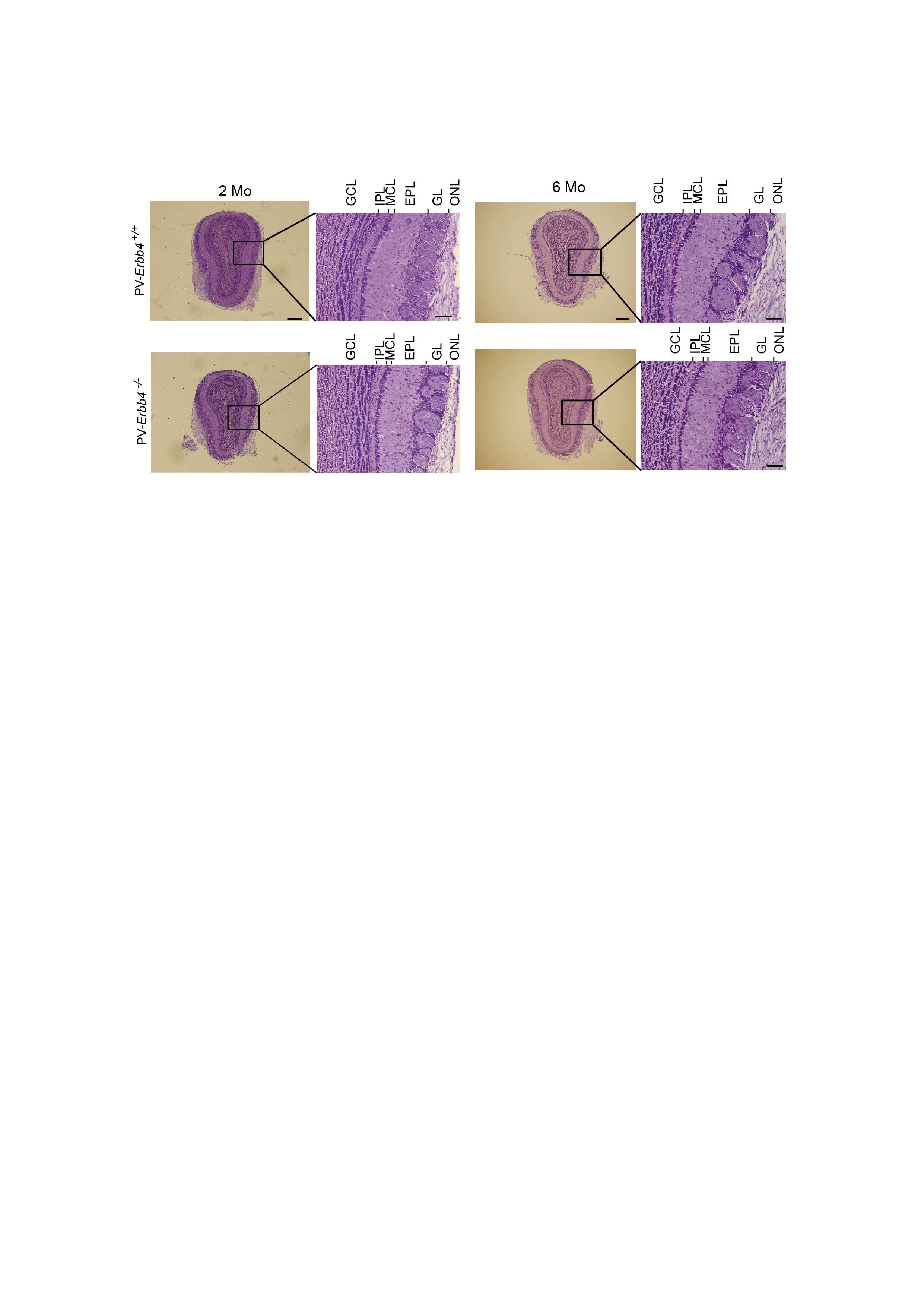

### Figure4 and supplement figure 1

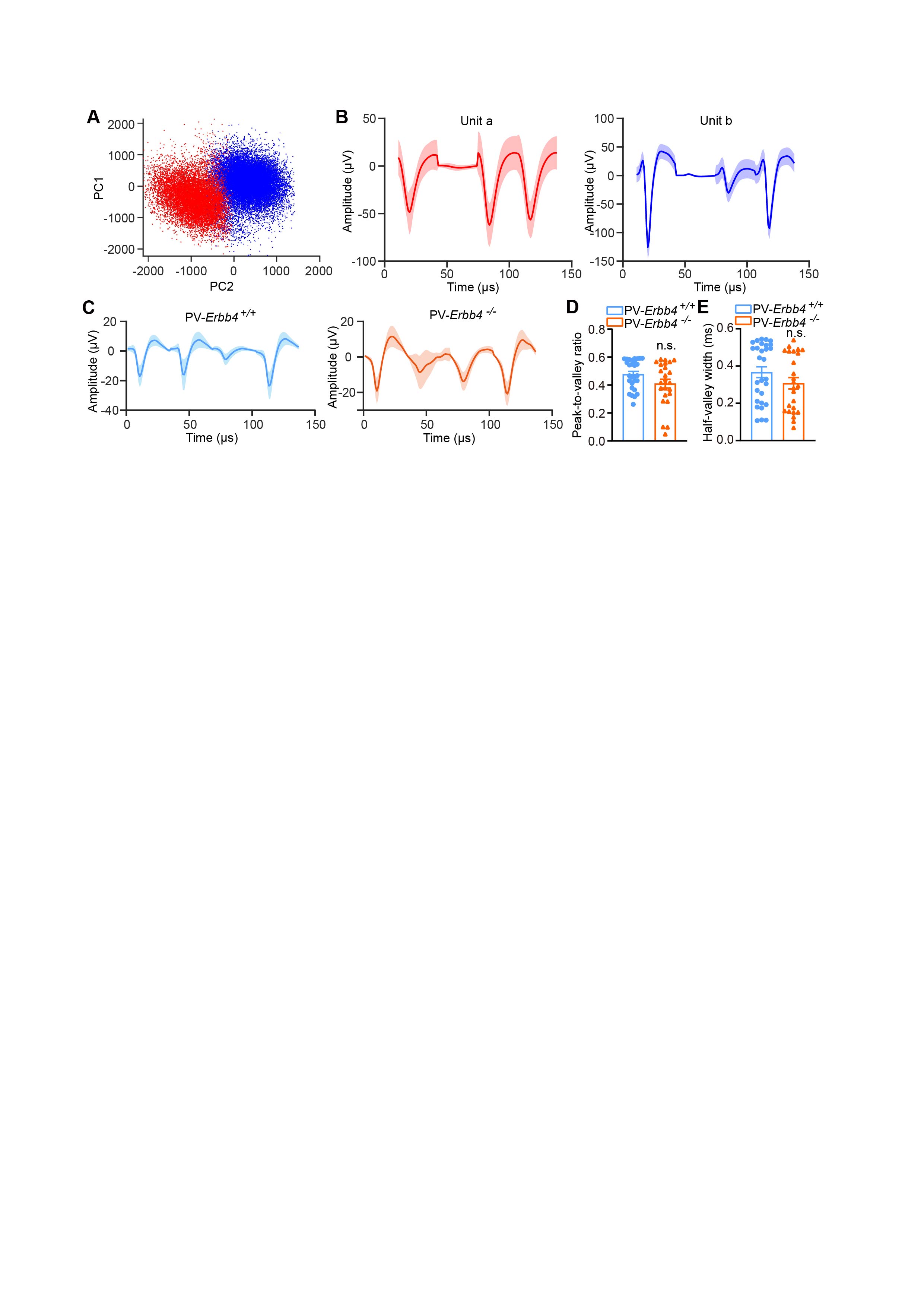

### Figure 7, Supplementary Figure 1

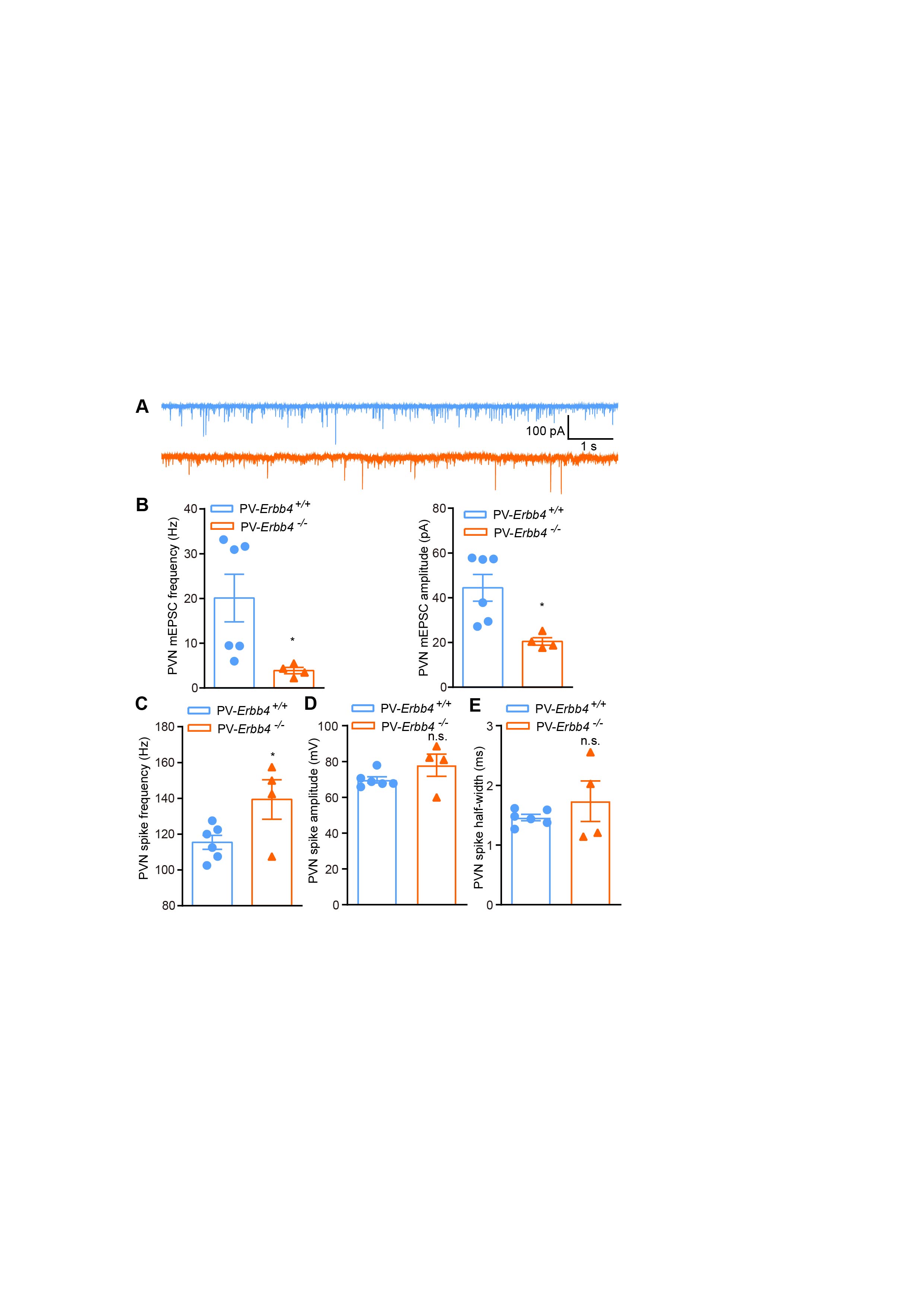

### Figure 8 and Supplementary Figure 1

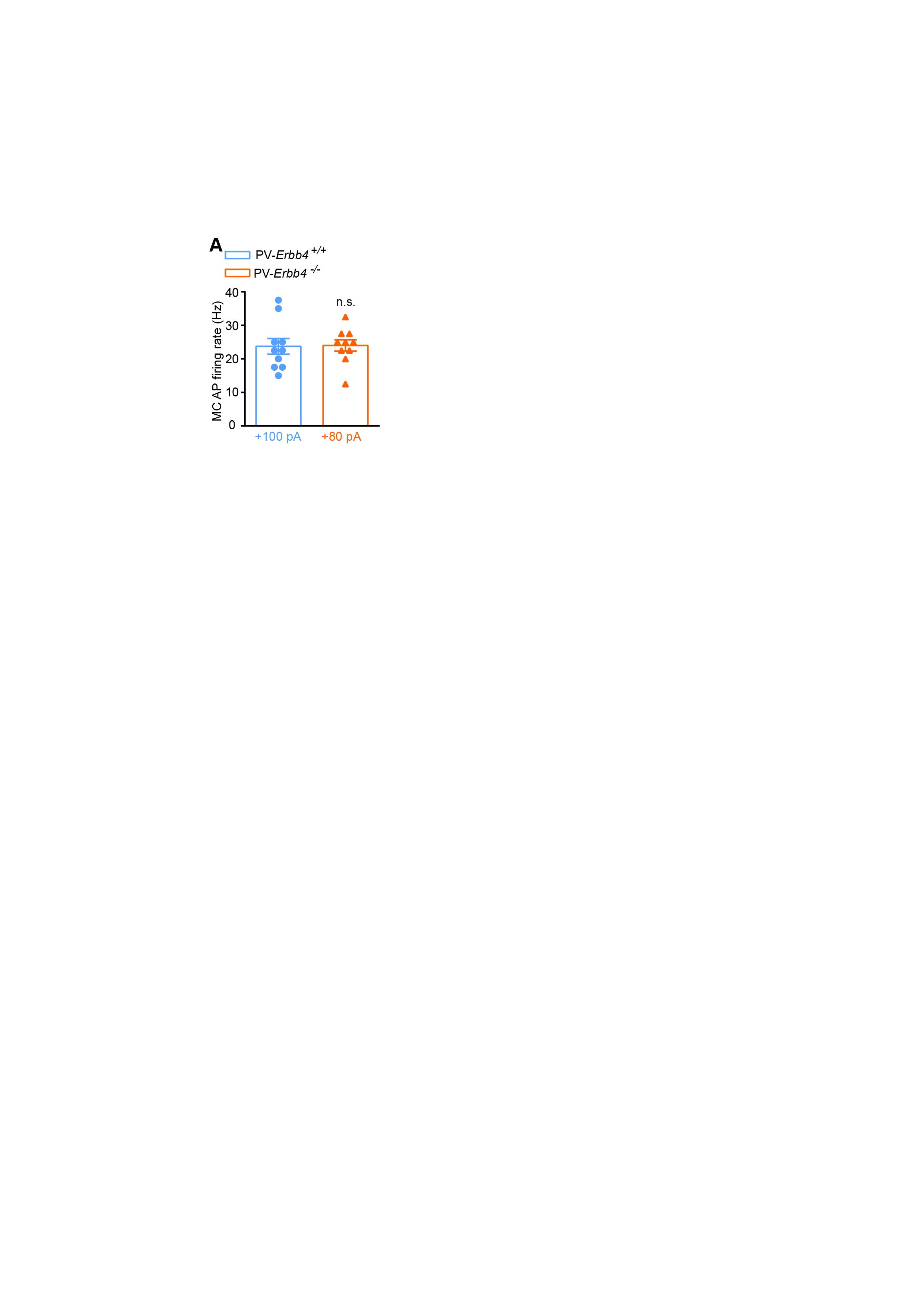

### Figure 9, Supplementary Figure 1

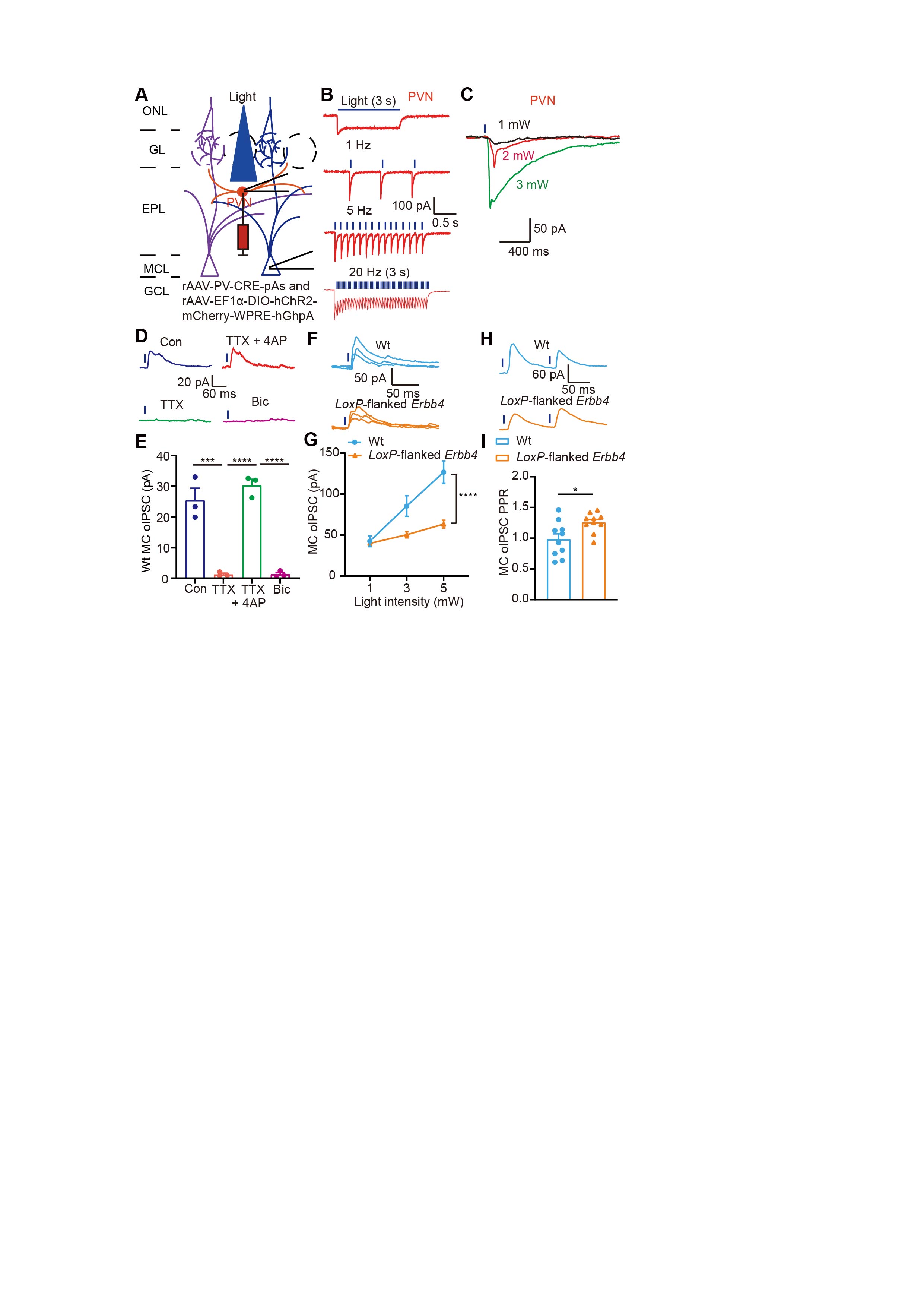
